# Staying in the club: Exploring criteria governing metacommunity membership for obligate symbionts under host–symbiont feedback

**DOI:** 10.1101/2020.01.10.901314

**Authors:** Vignesh Venkateswaran, Renee M. Borges

## Abstract

1. Metacommunity membership is influenced by habitat availability and trophic requirements. However, for multitrophic symbiont communities that are closely associated with host plants, symbionts and hosts may additionally influence each other affecting membership criteria in novel ways. For example, failure of beneficial services from a symbiont could elicit a response from the host that impacts the entire community. Understanding such host–symbiont feedback effects on symbiont community membership can be crucial for understanding symbiont community structure and function.
2. We investigate membership for a multitrophic insect symbiont community where symbionts colonize host inflorescences during specific developmental stages termed colonization windows. Inflorescences are host-derived organs and serve as habitat microcosms. Symbionts exhibit a diversity of interactions ranging from mutualism to parasitism. Hosts exhibit immediate feedback by aborting inflorescences not pollinated by mutualistic symbionts and habitats are consequently lost for all other symbiont species. Using relevant empirically measured microcosm parameters, we simulate symbiont dispersal from and colonization of other host inflorescences. We vary host densities and symbiont colonization window lengths, and track the persistence of each symbiont species in the metacommunity based on the temporal availability of the resource and the trophic position of the symbiont.
3. Since the persistence of the microcosm habitat is dictated by pollination performed by the mutualist, the mutualist fares better than all other symbionts. For prey, the length of colonization windows was positively related with colonization success and symbiont persistence. For predators, the cumulative length of the colonization windows of their prey dictated their success; diet breadth or prey colonization success did not influence the persistence of predators. Predators also had a greater host-plant density requirement than prey for persistence in the community. These results offer valuable insights into host density requirements for maintaining symbionts, and have implications for multitrophic symbiont community stability.
4. *Synthesis*. Factors influencing symbiont community membership can be unique when host–symbiont feedback impacts host microcosm development. Special constraints can govern symbiont community membership, function and structure and symbiont persistence in such metacommunities.

## 1. INTRODUCTION

Local and global processes in metacommunities govern species persistence and overall community membership. Within local communities, competition and predation can influence persistence (Holt & Bonsall, 2017), while prey availability can affect predator membership (Holt, 2009). Additionally, global processes such as the ability to successfully disperse across habitats can impact local community membership as well as metacommunity membership (Leibold et al., 2004). For symbionts in obligate host–symbiont structured communities, symbiont-mediated effects on host fitness can feed back onto the persistence of the symbiont metacommunity itself (Miller, Svanbäck, & Bohannan, 2018).

When host and symbionts are jointly responsible for the development or persistence of the host or host organs in which the symbiont community assembles, the survival of either entity becomes paramount for community membership of all symbionts. Therefore, when host– symbiont feedback dictates the ontogeny of such organs, incorporating these feedback processes into symbiont community membership is vital. Such investigations are nascent (Miller, Svanbäck, & Bohannan, 2018).

Metacommunities can consist of distinct habitats that occur discretely in space and time (Leibold et al., 2004) with community members residing on and dispersing across habitats.

Similarly, in symbiotic metacommunities, each host may be likened to a habitat (or a microcosm) and the symbionts it harbors to species residing within that habitat. Hosts must be sufficiently abundant in space and time to enable successful dispersal and colonization by symbionts (Arneberg, Skorping, Grenfell, & Read, 1998; Venkateswaran, Shrivastava, Kumble, & Borges, 2017; Venkateswaran, Kumble, & Borges, 2018). But host numbers themselves may be mediated by symbiont effects on the hosts. In cases where host–symbiont interactions are characterized by obligate mutualistic dependencies (as they are in many symbiont communities), the survival of host and mutualistic symbionts is a pre-requisite for the persistence of the entire symbiont metacommunity (Figure 1a); e.g. in human microbiomes the core microbiome may be considered indispensable for human health and survival (Bäckhed et al., 2012), potentially making its persistence critical for other “non-core” microbial symbionts.

**Figure 1.**
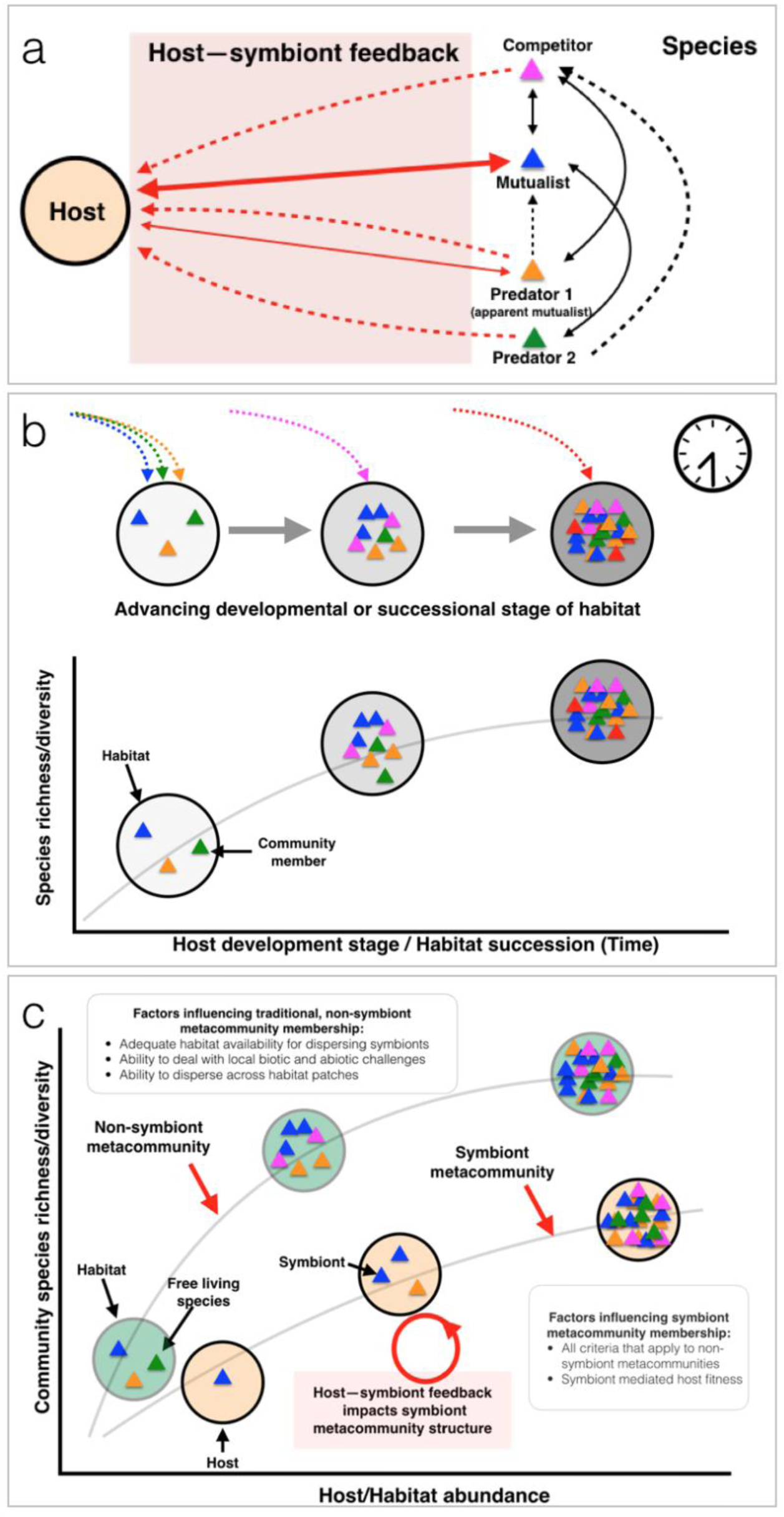
Rules dictating symbiont community membership. All species are depicted by triangles of different colors. Species are only symbionts where explicitly mentioned in the figure. **a**. Symbiont metacommunities can be fundamentally different from traditional communities consisting of free-living species assembling in habitat patches. Depicted is a hypothetical network of interacting symbionts and a host, all influencing the fitness of the host directly (solid arrows) or indirectly (dashed arrows). Each symbiont affects the fitness of the host and each other (signs indicate the impact on fitness). The red arrows exclusively indicate fitness effects on the host by symbionts. Specific relationships between hosts, parasites (of symbionts) and mutualists all serve to impact host fitness and eventually host abundance. Symbiont communities structured around a core mutualism (as indicated by the thicker red arrow) are likely influenced by the performance of the mutualism; the demise of either mutualist, or host sanctioning to punish cheaters may influence fitness of all symbiont species. **b**. The developmental stage or the successional stage of a habitat can restrict community membership, i.e. certain members may only be able to colonize a habitat at particular time durations when new, suitable niches are made available. For example, predators may only occupy a community after the colonization of prey. Such subsequent and conditional colonization may generally influence average local species richness, though not always in a positive manner as shown in Fig 1a. **c**. A hypothetical comparison between a traditional and a symbiont metacommunity. All else being equal, if symbiont metacommunities have more membership-limiting criteria governed by host–symbiont feedback, they may support fewer species with increasing host abundance and will exhibit species accumulation trajectories that are markedly different from those of traditional metacommunities.

Since the colonization of hosts and emigration from hosts may occur only during specific periods of host development (Mcfall-Ngai, 1994; Kitching, 2001; Srivastava et al., 2004; O’Neill, 2016; Borges, 2017), host ontogeny can heavily influence symbiont community membership. Therefore, the accumulation and release of symbionts may be contingent on the host’s own development as well as the number of other hosts in the population that release symbionts (Figure 1b) leading to two logical insights. First, when hosts develop asynchronously with each other in ecological timescales, they can facilitate the transfer of symbionts across them. Second, smaller time windows for colonization during host development should decrease host availability and symbiont colonization success (Venkateswaran et al., 2017; Appendix 1).

Even though the stable functioning and persistence of host and symbiont metacommunities are of considerable theoretical interest (Bordenstein & Theis, 2015; Douglas & Werren, 2016; Roughgarden, Gilbert, Rosenberg, Zilber-Rosenberg & Lloyd, 2018), membership criteria involving temporal aspects of colonization by symbionts, inter-symbiont interactions, and host–symbiont feedback have received little attention. Symbiont species accumulation and resulting symbiont biodiversity patterns may be markedly different owing to such host–symbiont feedback effects (Figure 1c).

In symbiont communities, such as gut-associated microbes and hosts, hosts usually outlive symbionts. In order to study feedback effects on symbionts, the long-term performance of hosts can be monitored. Alternatively, when hosts produce short-lived organs whose development depends on an assemblage of symbionts and whose ontogeny is fundamentally linked to host fitness and also demonstrates feedback effects, then investigation of such organs that serve effectively as habitat microcosms are powerful model systems to address such questions. We describe and investigate a multitrophic symbiont community of fig wasps that assembles in the inflorescences of a host plant, and that exhibits immediate host–symbiont feedback effects. Host plants abort inflorescences that have not received pollination services from a symbiont leading to habitat loss for all symbionts. We examine how metacommunity membership of multitrophic symbiont fig wasps associated with a single host plant species is influenced by host–symbiont feedback by simulating hypothetical and real fig wasp communities using relevant empirically measured community parameters. We vary three important variables: 1) host abundance (increasing number of trees), 2) lengths of the colonization window (equivalent to the oviposition window) for herbivorous and predatory symbionts, and 3) prey availability for predators. We further show how symbiont membership can be influenced by host–symbiont feedback and discuss implications for symbiont community composition and persistence in such ephemeral host-derived microcosms.

## 2. METHODS

### 2.1 Fig microcosms and associated symbiont communities, natural history and membership constraints

Figs and their fig wasp symbionts represent a model plant–insect symbiont community. Multitrophic fig wasp communities that occupy and develop within enclosed fig inflorescences are species-poor, consisting of no more than 30 species (Compton & Hawkins, 1992), but are characterized by rich interactions between symbionts and the host (Compton & Hawkins, 1992, Ghara & Borges, 2010). Such symbiont insect communities consist of obligate pollinating mutualists, competitors, and predators. Each symbiont confers a net positive, negative or neutral benefit on the host (Ghara & Borges, 2010; Segar, Pereira, Compton, & Cook, 2013; Krishnan & Borges 2014, 2015; Venkateswaran et al. 2017, 2018).

Each of the 800+ fig (*Ficus*) species produces many inflorescences (also called syconia; closed urn-shaped microcosms, singular=syconium), each of which is a host organ that represents an ephemeral microcosm (Cook & Lopez-Vaamonde, 2001). Wasps disperse from the natal inflorescence to another after development to adulthood (Figure 2) (Janzen 1979).

**Figure 2.**
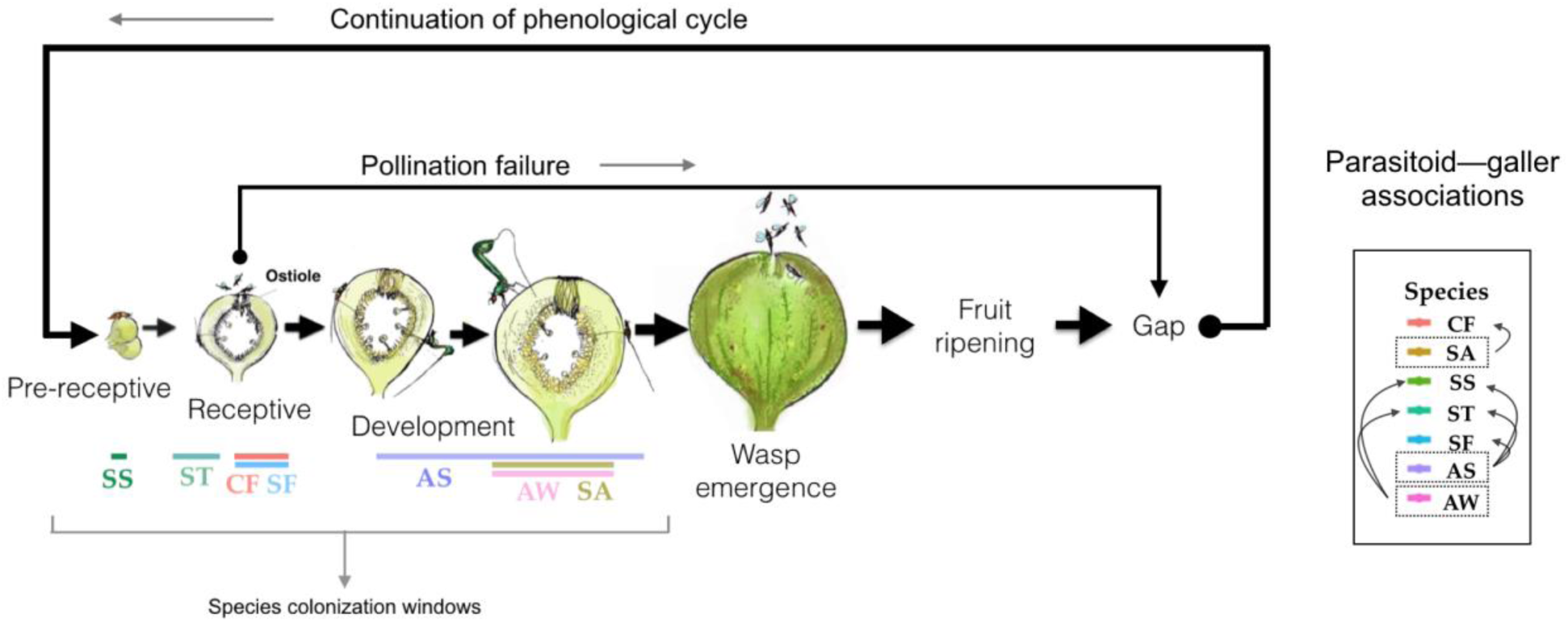
Ontogeny of the floral microcosm (i.e. syconium) in a typical fig with typical development phases. Since trees usually bear all their inflorescences in the same developmental phase, the phenology of a single inflorescence or microcosm represents the fruiting phenology of a tree. Pre-receptive = microcosm phase prior to pollen receipt; receptive = phase of pollen receipt; development = phase when wasps and seeds mature. Details of symbiont species are provided for the model cluster fig *Ficus racemosa*. The timing of arrival (oviposition window) of each wasp species during inflorescence development is indicated by colored bars under the different stages of development of the inflorescence. Species functional classifications and abbreviations are the following. Mutualist galler (herbivore): CF=*Ceratosolen fusciceps*. Gallers (herbivores): SS=*Sycophaga stratheni*, ST=*Sycophaga testacea*, SF=*Sycophaga fusca*. Parasitoids (predators): AS=*Apocrypta* species 2, AW=*Apocrypta westwoodi*, SA=*Sycophaga agraensis*. Parasitoids are presented within boxes and the parasitoid–galler associations are indicated by arrows.

Pollinating wasps induce galls within the inflorescence tissue, within which their offspring develop. Competing wasp symbionts also induce galls (herbivorous gallers) in the inflorescence and compete for oviposition space. Both pollinators and competing gallers feed directly from plant tissue within their galls. Late-arriving parasitoids (predators) feed on developing galler offspring and show variable but species-specific diet breadths (Ghara & Borges, 2010). We use the terms parasitoids and predators interchangeably. A few fig wasp communities also include seed-eating wasps (Pereira, Teixeira, & Kjellberg, 2007), but they are not considered in this investigation.

Figs often abort un-pollinated inflorescences. Such abortions have often been seen as a host sanction strategy against mutualists that cheat and offer no pollination services (Jandér & Herre, 2010). Abortions may also serve to avoid the cost of bearing expensive floral tissue when pollinators are absent. Since abortion of inflorescences in the absence of pollination represents a situation where the lack of services offered by the mutualist results in an immediate termination of the habitat (inflorescence) for the entire symbiont community, it likely influences symbiont persistence. Though cheater pollinators or gallers may drive the development of inflorescences, we do not incorporate such effects in this early stage of our investigation. Figs exhibit within-tree reproductive synchrony, and in the absence of local availability of pollinators, a tree will abort all its inflorescences (see assumptions in Methods and associated references).

For more details on the natural history, trophic associations, symbiont influences on the mutualism, and resource occurrence in fig wasp communities, see Appendix S1 and Figure S1. Wasps leave their natal trees owing to phenological characteristics that preclude their re-colonization (see Supplementary material) and colonize other trees by ovipositing into inflorescences; the next generation of wasps develop within the inflorescence and disperses to colonize another tree. Each wasp has a specific time window, called the colonization window or oviposition window (OW), during the development of the inflorescence, when the inflorescence bears suitable oviposition sites (Figure 2).

We investigate membership criteria by simulating the phenological resource landscape for a model fig species *Ficus racemosa* and its associated wasp community of seven fig wasp species (Ghara & Borges, 2010) and also for other hypothetical fig wasp communities. All wasp symbionts are indicated in Figure 2 and are hereafter capitalized and italicized in the main text.

### 2.2 Estimating lengths of phenological phases from field data

We used the fruiting phenology census from an earlier study on *Ficus racemosa* (Krishnan *et al*., 2014) to inform our simulations and to derive biologically relevant parameters for hypothetical fig wasp communities. For more details, see Appendix 1. In all, we analyzed the developmental progression for 13,846 inflorescences, observed over multiple fruiting cycles spanning two years across 20 individual trees and recorded the lengths of 41,122 complete phases. Using these values, we calculated the length of each phase and its associated natural variance (Figure S2) and incorporated the value of the phase lengths in the simulations described below.

### 2.3 Agent-based simulation framework for host developmental progression and microcosm colonization

We used a framework that simulates the ontogeny of multiple asynchronous hosts with symbionts dispersing across hosts. Within these simulations we investigate how the timing of colonization during host ontogeny, prey requirements by predatory symbionts, and the role of the mutualistic symbiont in ensuring habitat persistence (through host–symbiont feedback) affect symbiont community success. All these effects are tested for different host abundances.

The simulation framework we adopted is similar to Bronstein *et al*. (1990) and Kameyama *et al*. (1999) but incorporates other essential features. The earlier simulations investigated mutualist performance alone without mutualist–host feedback. Our simulations include an entire multitrophic symbiont fig wasp community with host–symbiont feedback effects and predator–prey dependencies (details of the code in Appendix S1 and S2). As in previous investigations, we do not investigate population dynamics of the trees since multiple fig wasp generations can occur within a single host tree generation. Also, as in earlier studies, we do not address the population dynamics of wasps; our simulations keep track of symbiont species harbored by any individual host at every time step. We also do not explicitly test spatial distribution of trees; rather, we test symbiont community performance with increasing host abundance either by increasing the number of hosts or by increasing the lengths of the colonization window for symbionts.

Fig phenology was simulated by parametrizing the length and the variance of each phenophase of the inflorescence from a natural dataset. Each of the six phenological phase lengths (PR: pre-receptive, R: receptive, D: development, E: wasp emergence, G: gap, FR: fruit ripening; Figure 2) were parametrized based on a phenological census of *F. racemosa* trees. Over-dispersion (with distributions spilling into negative ranges) was prevalent for short phases, especially where variances were greater than the mean. Hence, we adopted a zero-truncated Poisson structure to select positive, non-zero integers from a discrete distribution. The length of each phase with its underlying natural distribution was used in our simulations for all host trees. In this way, unique sequences were generated at each fruiting cycle for every tree. To determine when symbionts could colonize hosts, symbiont colonization windows were restricted to fractions of the lengths of each phenophase. These values were specified based on specifications for each dummy model, or based on observational data for the simulation of the real community of *F. racemosa* (from Ghara and Borges 2010, see below). At the beginning of the simulations, every tree began at a random point in its generated developmental sequence. After a complete developmental cycle, a new and unique sequence was generated. If a tree was not colonized by a mutualist, the tree skipped the intervening D (development), E (wasp emergence) and FR (fruit-ripening) phases and entered the G (gap) phase. In this way abortions occurred only in the absence of pollinator services (Figure 2).

Symbionts were allowed to emerge and colonize trees. All symbionts were introduced by being allowed to emerge, disperse and colonize trees in the first round of release of each tree, after which the persistence of each symbiont species was contingent on its subsequent successful colonization. On each day, the trees that were releasing wasps and trees that were suitable for wasp colonization were noted. Additionally, the presence or absence of symbionts was recorded in each tree for each day. Each symbiont species successfully colonized a host provided that the timing of emigration of the symbiont matched the occurrence of a host at a suitable developmental phase on the same day. In this way, a tree acted as a suitable oviposition resource when its phenological phase corresponded with the timing of the OW of a particular symbiont (when it was suitable for colonization). For parasitoids (predators), an additional pre-requisite for colonization was that the tree needed to harbor the parasitoid-specific prey (galler species).

Each simulation was run with discrete tree population sizes ranging from 10 to 100 in increments of 10 for 1800 days (5 years); a previous investigation on fig tree–pollinating wasp stability incorporating the same agent-based framework revealed that a 5-year period satisfactorily predicted long-term (1000 year) outcomes (Anstett, Hossaert-McKey, & McKey, 1997). Colonization success was measured by two metrics: the total number of colonization opportunities (CO), i.e. the number of times trees were available for colonization when a symbiont was released, and the proportion colonization success (PCS) for each wasp symbiont species, i.e. the ratio of the number of times wasps were successfully able to colonize a resource to the total number of times they were released in the simulation. The two metrics are important because they provide unique measures of performance of each symbiont. PCS indicates the probability that a symbiont will successfully colonize a resource in the landscape when released by a host. However, the frequency of symbiont release could still be very low and curtailed by other factors. CO, on the other hand, also captures the frequency of the release of the symbiont. Therefore, these metrics are distinct and are useful when considered together.

Thirty iterations were conducted for each host density tested. We do not account for space in our model and use the term tree abundance to indicate the number of potentially accessible hosts (host density).

### Assumptions

1. All wasp species live only for a single day and they can colonize trees only when their release coincides with the availability of a suitable plant host on the same day. Though wasps can live more than a day when provided with ad-libitum water and sugar supply (Ghara & Borges, 2010), studies also suggest that wasps during flight are susceptible to extreme exhaustion resulting in reduced lifespans of less than a day (Jevanandam, Goh, & Corlett, 2013; Venkateswaran et al., 2017). Extrinsic mortality can also be high due to predation pressure, desiccation, and high temperatures (Ranganathan & Borges, 2009; Ghara & Borges, 2010; Jevanandam, Goh, & Corlett, 2013), further reducing the longevity of wasps.
2. When no colonization by the mutualist occurs during the receptive phase (pollination failure), all inflorescences in the tree are aborted due to the lack of pollination and the tree enters the gap phase after which the tree initiates another cycle (Figure 2), making habitat resources unavailable for symbionts on the tree for that cycle.
3. Successful parasitoid (i.e. predator) colonization has no effect on subsequent prey (galler) release; i.e. we do not account for predator–prey dynamics. Although such predatory effects can also influence symbiont membership, their inclusion demands a separate investigation which is beyond the scope of this study.

### 2.4 Features of the simulation model variants

We used four dummy models (and a dummy model variant) to understand the functioning of the real fig wasp community of *F. racemosa* (Figure 3). All parameter values used in each dummy model and in the real model are listed in Table 1. All effects are compared as a function of host abundance (Appendix S1). In the dummy models, we consider the performance of representative wasp species as functions of their OWs and diets. All four dummy models included the mutualist colonization and tree abortion events. Sensitivity analysis were conducted to ensure that parameter changes had a robust effect on all colonization success output values despite the inherent stochasticity present in each model (Appendix 1).

**Figure 3.**
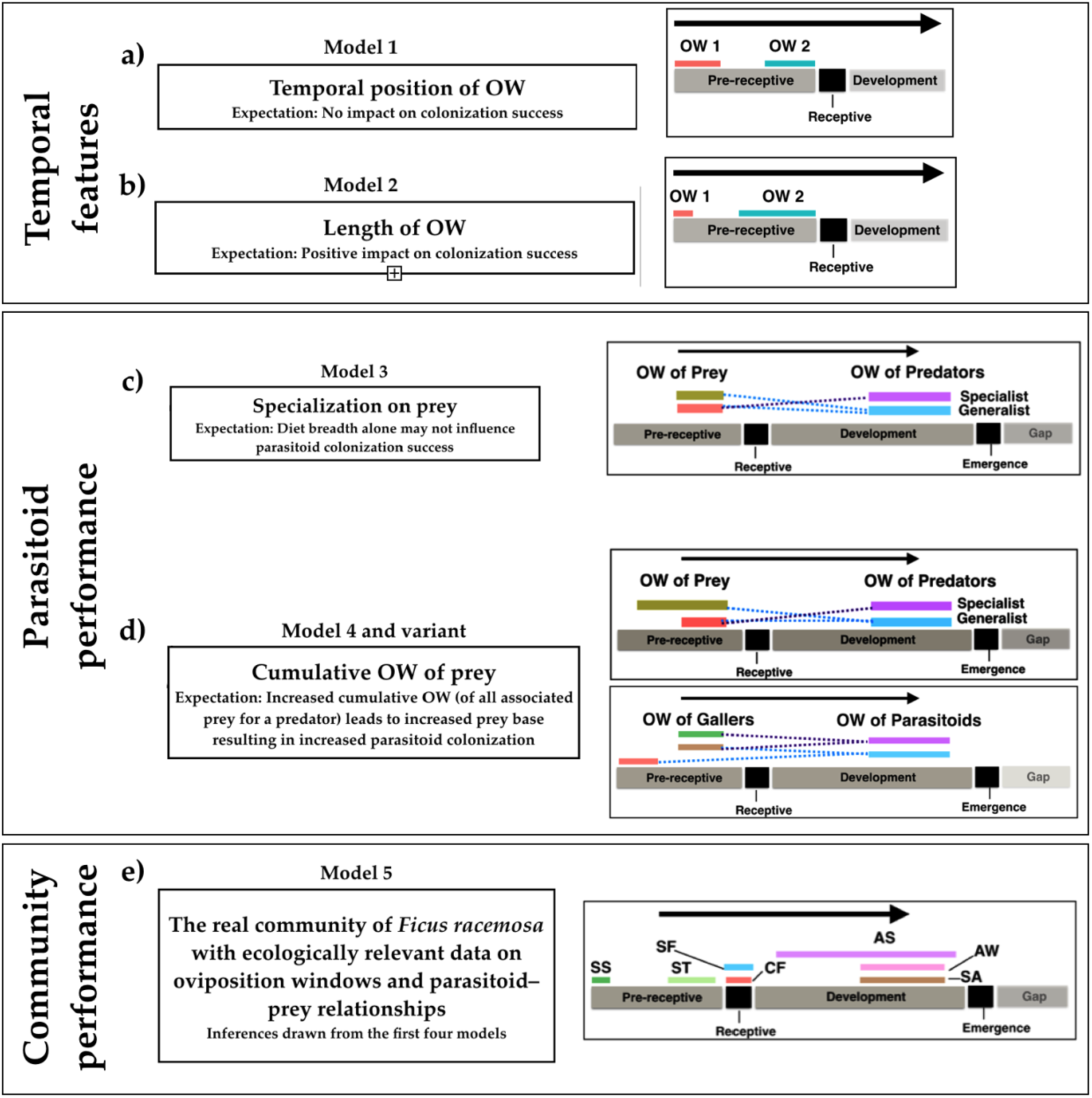
Description of the models. **a)** The influence of colonization success when OWs occur at different times of development of the microcosm. **b)** The influence of colonization success when OWs are of different lengths. **c)** The influence of the number of prey species (prey specialization) on colonization success of two hypothetical parasitoid (predator) species. **d)** The influence of prey colonization success on colonization success of parasitoids (also see Appendix S1 for variant of Model 4). Models 3 and 4 are applicable to parasitoids in the community. **e)** The colonization success of members of the real *F. racemosa* wasp community (species abbreviations are as indicated in Figure 1). For parameter details see Table 1.

**Table 1.**
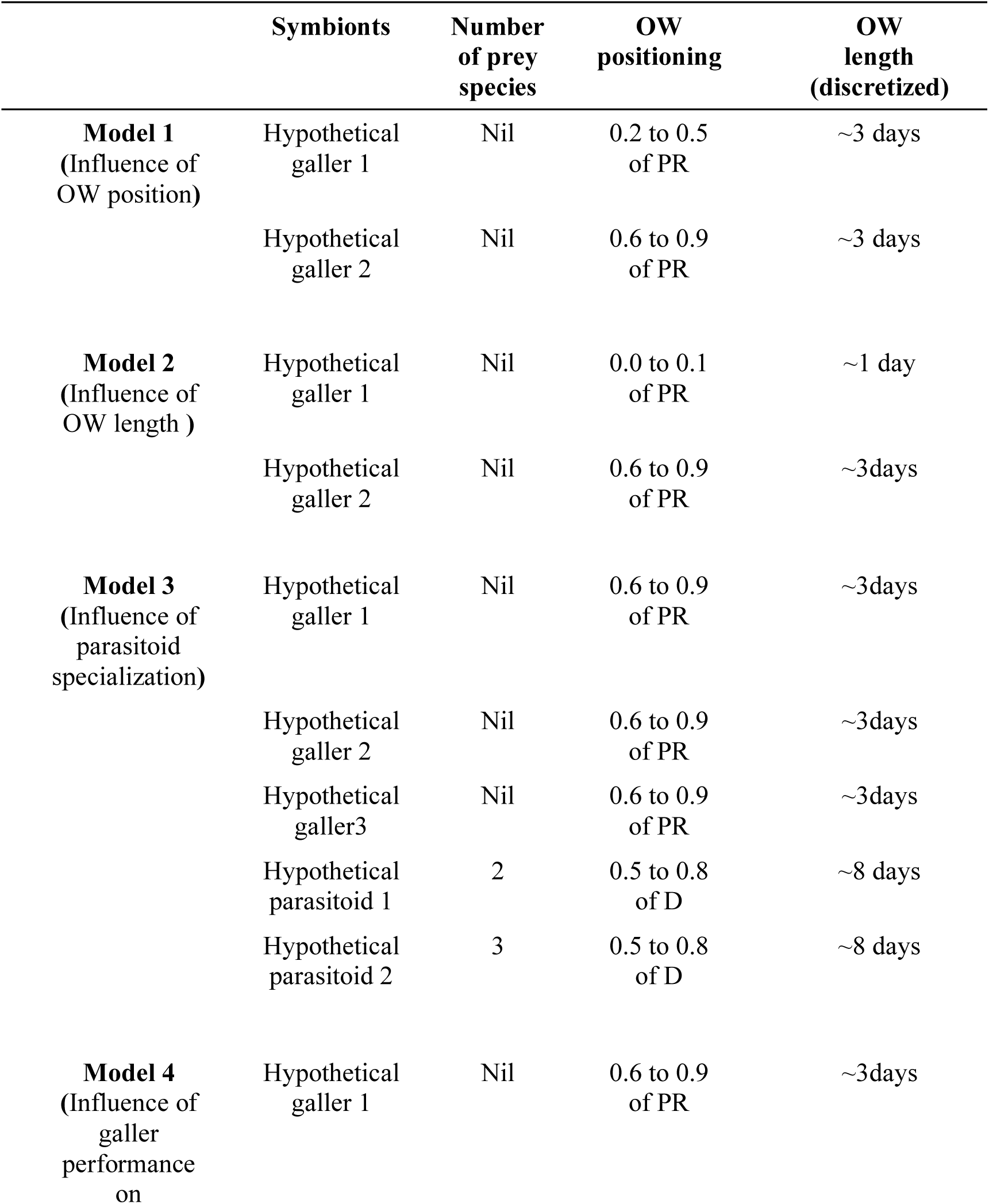
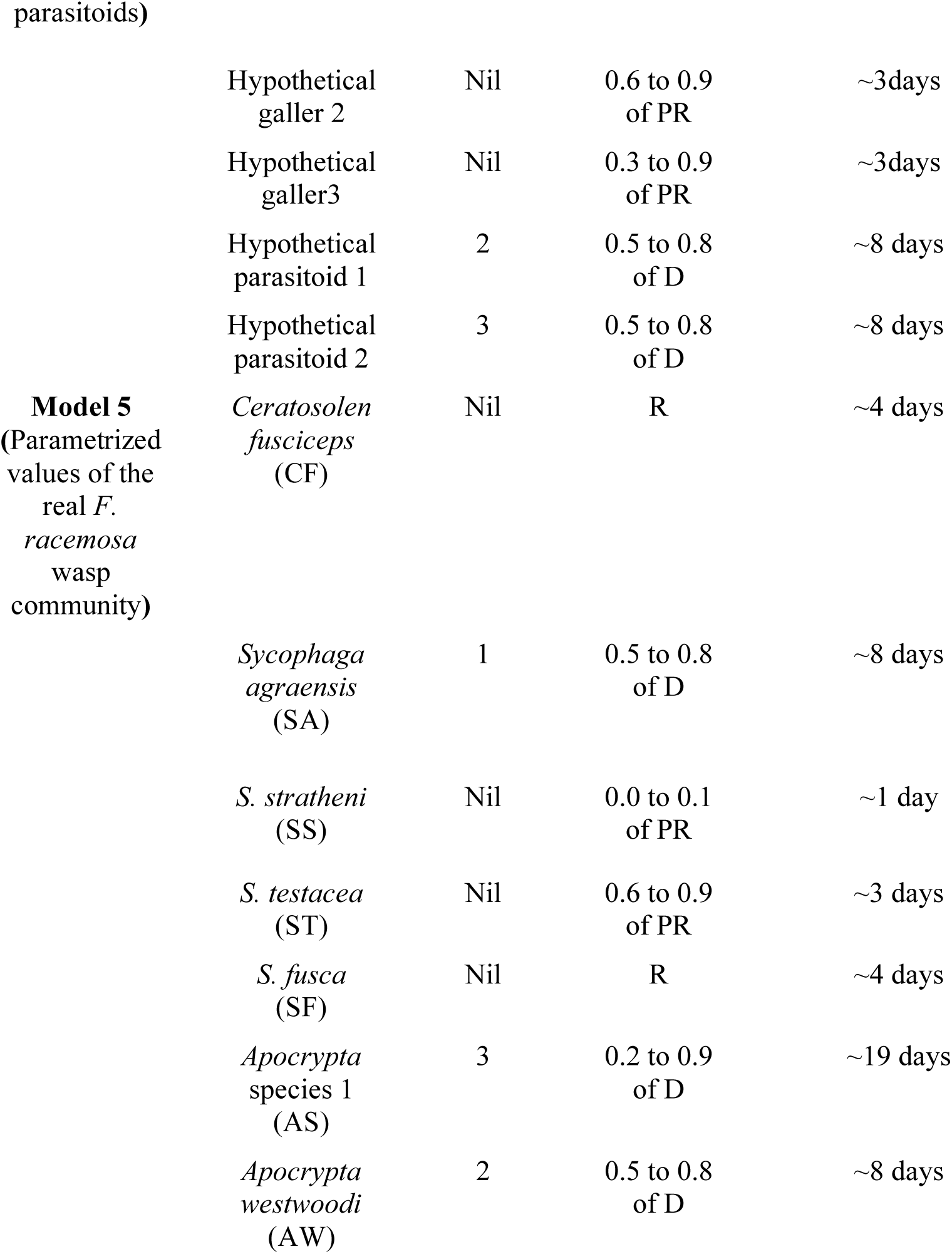
The positions and lengths of oviposition windows (OWs) of the different wasp symbionts and the number of hosts (for parasitoids) in each model. The abbreviations indicate the developmental stage of the inflorescence: PR=Pre-receptive phase, R=receptive phase, D=Developmental phase. Wasp abbreviations for Model 5 are the same as in Figure 1.

In Model 1, we vary the position of the OW (Figure 3). As both changed positions are within the pre-receptive window, and there is no inflorescence abortion that occurs within the pre-receptive window, we expect the OW position to have no effect on CO (Figure 3, Figure 4). In Model 2, we vary the length of the OWs (Figure 3). Since a tree is colonized by a wasp if and only if there is another tree releasing this wasp species within its OW, therefore, the likelihood of colonization should increase with OW length (Figure 3). We also note from our preliminary analyses (results not shown) that if the OW of two species completely overlap, then the trees holding them after colonization are also identical. This is because in our simulations, if release and colonization events coincide, then colonization happens with probability 1. Hence there is no difference in the colonization events between the two species.

**Figure 4.**
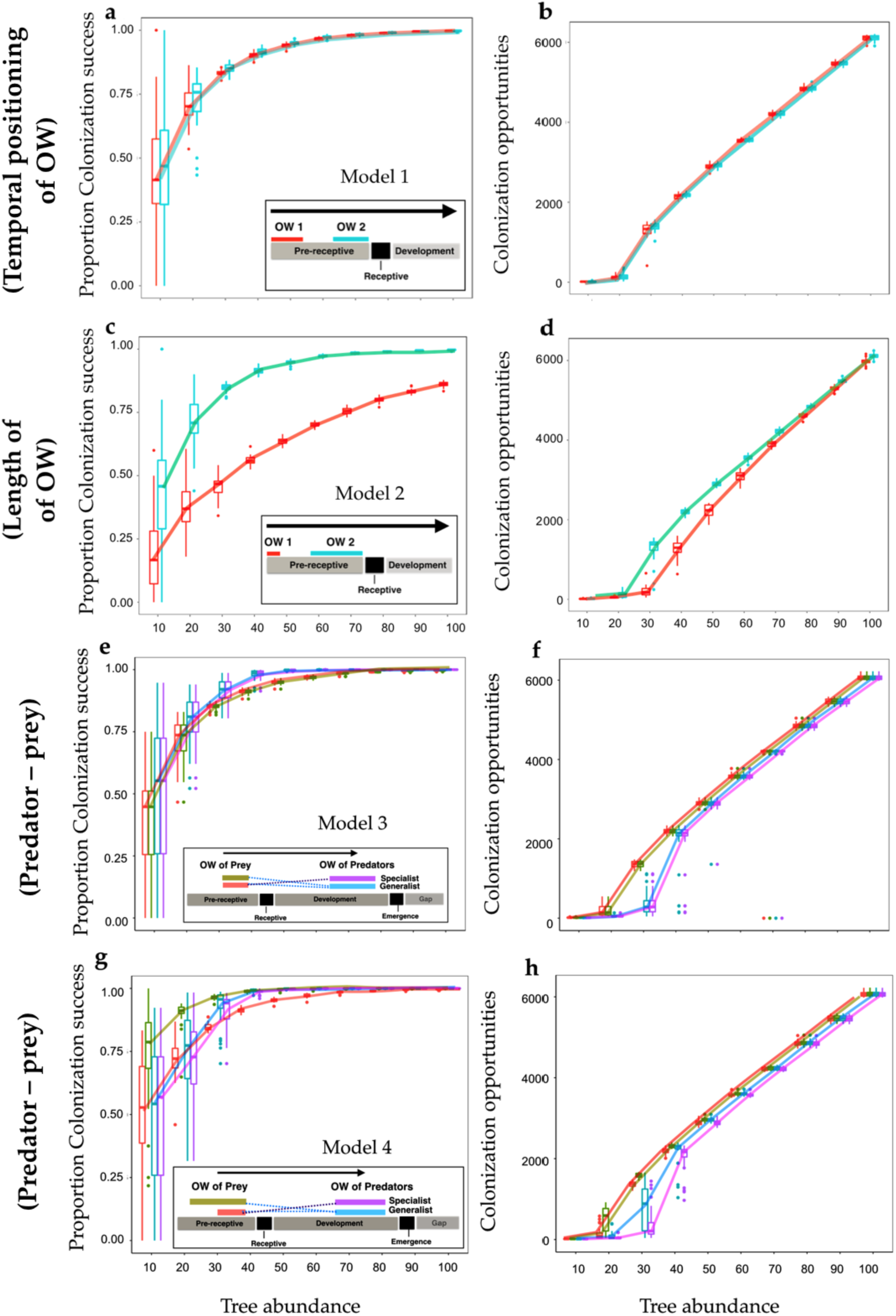
The colonization success (PCS and CO) of predator and prey as a function of tree abundance. The length and positioning of the oviposition windows of each wasp species over inflorescence development are indicated in the in-laid schematics. **a)** Model 1: Two hypothetical galler wasps that have the same OW length but that differ in their colonization sequence. The sequence does not affect colonization success. **b)** Model 2: Two hypothetical galler wasps that have differing OW lengths. Longer OWs enhance colonization success. **c)** Model 3: Two hypothetical parasitoids (i.e. predators), one specializing on two hypothetical galler prey species (broad diet breadth) and another specializing on one prey types (narrow diet breadth). **d)** Model 4: The same as Model 3 except that the galler that is exclusively preyed upon by the generalist predator has a longer oviposition window than the other two gallers; this indicates that the subset of trees with more prey in general enhances parasitoid colonization rates (also see the variant of Model 4 in Fig S3). All models incorporate mutualist–host feedback; the performance of the mutualist is not shown in these models.

In Models 3 and 4, we consider the CO of parasitoids, and their dependence on the prey species. In Model 3, we compare the CO of two parasitoids— one that feeds on two prey types (generalist), and the other that feed on only one prey type (specialist), such that the OWs of the two prey coincide (Figure 3, Figure 4), as do that of the predators. A priori, we may expect that the generalist parasitoid does better, as it has more prey options. However, we also note that because the prey OWs coincide, all the trees holding one prey type also hold the other in our simulations. Hence there may be no difference in the success of these two parasitoids. In Model 4, we again compare the CO of generalist and specialist parasitoids, but now the OWs of the two prey are not the same because they are of different lengths and have different positioning (Figure 3). We now expect the generalist parasitoid to do better for two potential reasons: 1) feeding on a prey that has greater persistence (with larger OW) may itself enable the predator to perform better since larger windows increase the number of trees harboring the prey and will aid predator colonization or, 2) an increased number of trees holding all relevant prey because of decreased overlap of prey OWs can also allow for better predator performance. Both of these factors could lead to an increased prey base and requires to be tested independently.

We use a variant of Model 4 to disentangle the role of better performance of prey because of increased OW vs. the role of an increased non-overlapping cumulative oviposition window. Here, we address how the extent of overlap of prey OWs dictates parasitoid success while keeping the colonization success of all prey constant. Unlike model 4, in the Model 4 variant, we incorporate two predators and three prey (Figure 3). Each predator specializes on two prey, (with one shared prey species). All prey species have equal OW lengths (Figure 3d). For one of the predators, the two prey have completely overlapping OWs while for the other predator, the two prey have completely non-overlapping OWs. We expect that the predator with prey having overlapping OWs would perform worse—the trees holding each prey species would be the same, all else being equal.

Lastly, we simulated the real model with parameter values and species associations of the fig *F. racemosa* and its wasp community (Model 5=the real community) (Figure 3). This model was run to obtain a priori expectations for how the real wasp community may function to guide future studies. Results of this model were interpreted by referring to the results of the previous models.

All simulations were run using R 3.3.1 (R core team, 2017). The code for all model variants can be found in Appendix S2. For Models 1–4, Mann-Whitney-Wilcoxon tests were conducted to examine differences in the performance (CO and PCS distributions) of the two hypothetical wasps (gallers or parasitoids) at every tree abundance regime. Dunn tests with Bonferroni corrections at an alpha value of 0.05 were also conducted for the species in the simulation reflecting the real *F. racemosa* community (Model 5) at every tree abundance regime.

### 2.5 The influence of tree abundance on extinction probabilities and symbiont persistence

To understand the influence of host availability on persistence of the symbiont community, we also calculated the extinction probability of each symbiont in the real community for the duration of our simulations, i.e. the number of times a species went extinct in the 30 iterations for differing tree abundances (from Model 5). We investigated extinction probability as a function of tree abundance.

## 3. RESULTS

### 3.1 Influence of host–symbiont feedback (microcosm persistence) on community structure

Our results show that host densities and symbiont colonization windows affect species persistence or extinction. The length of each phase and associated variances estimated from the natural data set are presented in Figure S2. Higher tree abundance resulted in successful colonization of *CF* (pollinator), which allowed microcosms to persist (Figure S4). With only few trees, abortions were common (Figure S4a) which decreased with increasing tree abundance (30 trees, Figure S4b; 100 trees, Figure S4c).

## 3.2 Model results

### 3.2.1 Models 1 and 2

Symbiont oviposition window (OW) temporal sequence (Model 1) did not affect either the proportion colonization success (PCS) or colonization opportunities (CO) (Figure 4a, b; Table S2) unlike the length of the OW (Model 2) (Figure 4c, d; Table S2). Longer durations of resources aided in colonization of symbionts as expected. Colonization was inferred using PCS and CO; there were no significant differences between PCS or CO for the two galler species in Model 1 while there were significant differences in PCS and CO in all tree abundance regimes in Model 2. PCS, more than CO, increased with increasing tree abundance in all simulations for all symbionts (Figure 4c, d; Table S2; Table S3a, b). These results were confirmed by our sensitivity analysis showing that our models were robust despite the stochasticity (Appendix 1).

### 3.2.3 Models 3 and 4

As expected initially (Figure 3), more prey types did not necessarily increase predator performance. The results of Model 3 indicated that predator diet breadth alone did not affect colonization in our simulations (Figure 4e, f; Table S2).

The increased colonization opportunities for parasitoids through increased galler OW length in the results of Model 4 (Figure 4g, h) were as expected (see both PCS and CO of gallers and parasitoids). We tested the effect of staggering prey OWs without increasing their lengths on predators through the variant of Model 4 (Model 4b, Figure S3). This model revealed that, for parasitoids with the same number of galler prey species, those feeding on gallers with staggered OWs had greater colonization success (Model 4b, Figure S3). This result was again confirmed by our sensitivity analysis (see Appendix 1).

These results collectively indicated that the cumulative length of non-overlapping OWs of prey types was positively related to parasitoid colonization. A high cumulative OW length could be achieved by either a single prey type with long OW or multiple prey types with non-overlapping windows that add up. This cumulative length of prey OWs was necessary and sufficient to explain how all prey species attacked by a predator positively influenced predator. Diet breadth and prey colonization success alone were not satisfactory at explaining colonization performance of predators.

### 3.2.5 Model 5 (the real *F. racemosa* symbiont community)

The galler *SS* had the lowest PCS and CO at all tree regimes because *SS* had the smallest OW length (Figure 5a, b; Table 1). The pollinator *CF* had the highest CO (Figure 5b). At low tree abundance, *CF* (mutualist) populations crashed leading to the extinction of all other wasp species (Figure S5 a–e). The parasitoids *AW* and *SA* had the same OW lengths (∼ 8 days) and they showed no difference in their PCS with respect to each other at all tree regimes despite specializing on different prey types (Table 1, Figure 5a); this result was consistent with that of Model 3. However, *SA* had a higher *CO* than *AW* since it parasitized *CF* that was more successful than any of *AW*’s prey (Figure 5b), a result that was consistent with Model 4. *ST* had an OW between that of *SS* and the other five wasp species; its PCS curve showed a faster increase with increasing tree abundance as compared to *SS* but not when compared to the other wasp species (Figure 5a). *AS* with the highest OW (∼19 days, Table 1) showed the steepest rise in PCS with tree abundance (Figure 5a). *AS* also parasitized the most successful galler (*SF*) and, consistent with the results of Model 4, showed the highest CO compared to the other parasitoids (Figure 5b).s

**Figure 5.**
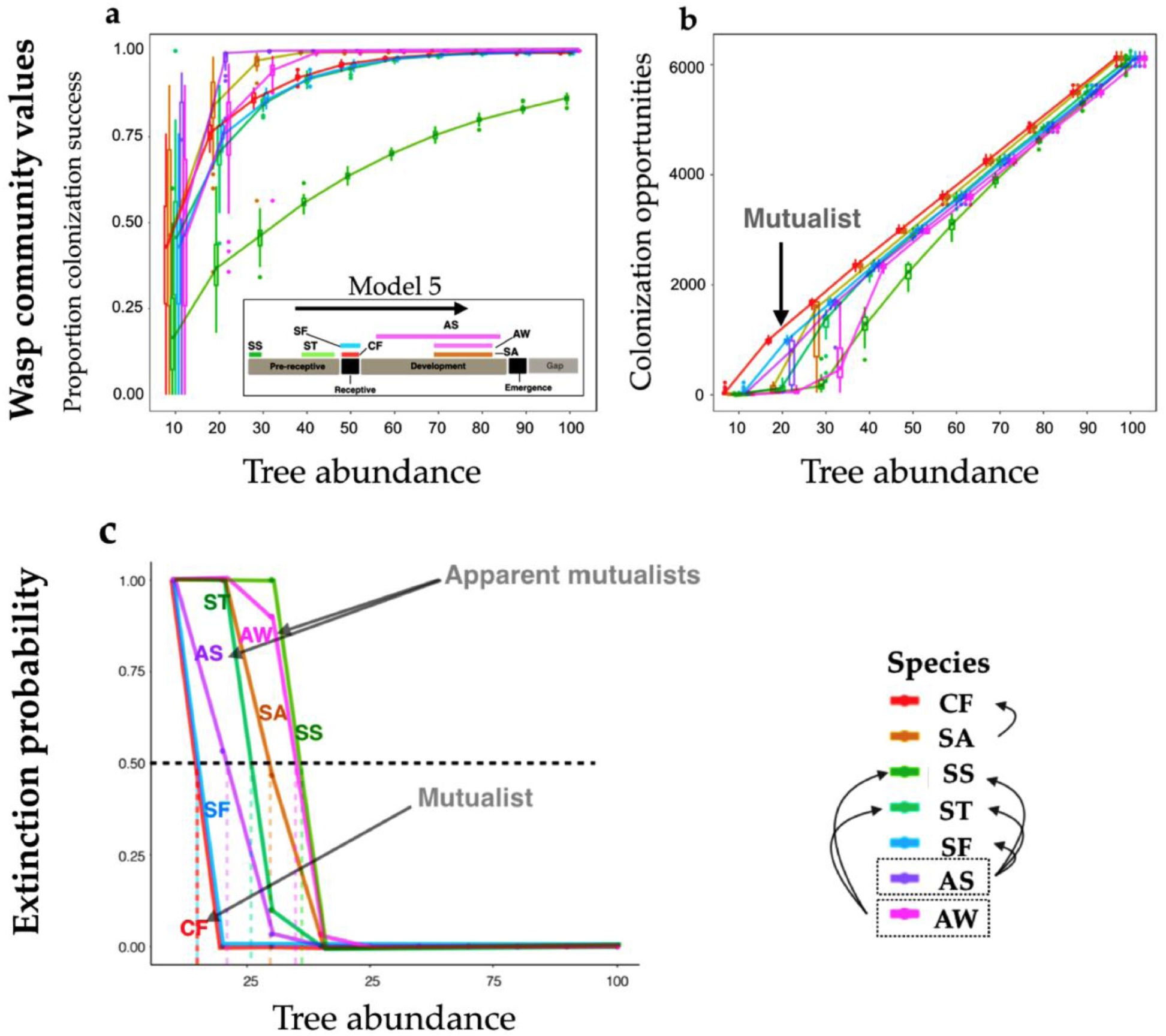
The colonization success (PCS (**a**) and CO (**b**)) for the real fig wasp community of *Ficus racemosa* as a function of tree abundance. Both proportion colonization success (PCS) and colonization opportunities (CO) values suggest that species with larger oviposition windows are more successful at colonizing hosts and parasitoid performance is based on prey (galler) availability, consistent with the previous models. However, the pollinators (*CF*) outperform all other wasps at all tree abundance as indicated by their COs. **c**) The probabilities of extinction with increasing tree abundance for each species in the real community of *F. racemosa*. The horizontal dotted line represents a 50% extinction probability. Vertical colored (dotted) lines indicate the tree abundance required to sustain the wasp species for five years with a 50% extinction probability. Predators (parasitoids) are indicated within dotted boxes and arrows indicate their galler prey. Species abbreviations are the same as in Fig 2. Predators (i.e. parasitoids) are indicated within dotted boxes and arrows indicate their galler prey.

In all models, the variability in PCS and CO for wasp species was lowest at high tree abundance (Figure 5 a,b) reflecting the decreased variability and an increased guarantee of resource occurrence with increasing tree abundance in the simulations. When OW values were the same (as for *AW* and *SA*; *SF* and *CF*), PCS values at all regimes were the same (statistically non-significant) while CO values were different (Table S4a, b), consistent with the results of Model 4.

### 3.3 The influence of tree abundance on extinction probabilities of species in the real *F. racemosa* symbiont community

The persistence of the mutualist was the highest at all host densities with no other symbiont species having higher survival probabilities. The minimum host requirements for persistence were highly dependent on the length of the colonization window; species with smaller OWs required more trees for persistence (Figure 5c). For parasitoids (i.e. predators), the minimum tree abundance required was never less than their most successful prey in the simulations.

Parasitoids, despite having larger OWs than the gallers (such as *AS*), were more likely to face lower absolute colonization success than gallers, due to their additional dependency on presence of galler prey within inflorescences. For instance, though the OWs of *SA* and *AW* (parasitoids) were of identical duration, the extinction probability of *AW* was greater than *SA* (as seen in tree regimes of 25 to 50, Figure 5c). The parasitoid *SA* is dependent on *CF*, which is the mutualist and performs better than any other wasp in any tree regime including *ST,* a galler prey species for *AW*. Therefore, the observed differences in extinction probabilities between the parasitoids *SA* and *AW* in the model indicated that they were contingent on the success of their specific gallers. These results provide a preliminary framework for future theoretical and empirical investigations.

## 4. DISCUSSION

Symbiont communities can be influenced by host development and host–symbiont feedback. In our investigations, such feedback ensured that no symbiont out-performed the mutualist (as inferred through the colonization opportunity (CO) measure, Fig 5b). Typically host–symbiont feedback effects are addressed with respect to responses between host and symbiont numbers (Mihaljevic, 2012). In fig symbiont metacommunities, symbiont wasps and host plants drastically differ in lifespan. Many inflorescence abortions may occur within the host’s lifespan without host mortality. Immediate host–symbiont feedback without host demise may be highly beneficial for metacommunity persistence since non-mutualist numbers may be immediately controlled relative to the pollinating mutualist performance, without the immediate demise of host or reduction in host numbers. With respect to host symbiont communities structured around a core mutualism, this predisposes the mutualist symbionts to greater stability and persistence at any host density, thereby increasing the persistence of the entire metacommunity. In the absence of feedback through abortions after mutualism failure, such stability will cease to exist.

The impact of such short-term effects without host demise and long-term effects, including an influence on host numbers, on the persistence and maintenance of symbiont metacommunities requires more attention. Such tests may even be performed in fig wasp systems in which parasitic galler wasps are able to hijack the host microcosm development and allow it to persist even in the absence of pollination (Krishnan *et al*., 2014; pers. observ.). Here, symbionts other than the pollinators may regulate host–symbiont feedback and fig wasp symbiont community structure. However, there has been no rigorous work on quantifying the effect of non-pollinating wasps in driving fig development even in the absence of pollination.

Mutualisms are often context dependent (Hoeksema & Bruna, 2015), and overexploitation by mutualists may reduce mutualist service, which may then transform into parasitism. Beside host sanctions to filter out such cheaters, mutualism stability may also be conferred by parasites or predators of the mutualists such as by parasitoids of the pollinator when pollinators over-exploit resources (Dunn et al., 2008). However, to prevent overexploitation by competitors or by the predators of the mutualist, other parasitoids in fig wasp systems may help confer stability by preying on non-mutualist species that directly compete with the mutualist (Yadav & Borges, 2017). Such parasitoids, therefore, act as apparent mutualists (Krishnan et al., 2015), and may also be very important for the stability of such mutualisms (Krishnan et al., 2015). Because our model reveals that such parasitoids that serve as apparent mutualists require a higher abundance of trees for their persistence than the mutualist, rethinking the minimum host abundance to account for the stability of beneficial symbionts in higher trophic levels is crucial and perhaps has valuable conservation implications (Shanahan, So, Compton, & Corlett, 2001). This is especially relevant since such apparent mutualists may confer “top-down” stability to the community (Estes, Tinker, Williams, & Doak, 1998). In many symbiont metacommunities, mutualists may be a prerequisite for community establishment and may be more persistent than other symbionts, while apparent mutualists may contribute to community stability. This requires an accurate characterization of trophic associations between symbionts; *F. racemosa* is perhaps the only fig wasp system where predator–prey relationships between gallers and parasitoids of the entire community have been established experimentally (Yadav & Borges, 2017).

Increasing host densities should support greater symbiont colonization and persistence as demonstrated by the results of all our models. However, our results also clearly show that the length of colonization window interacts with the development of habitats to eventually influence colonization success (as indicated through PCS and CO) and overall symbiont community persistence. We propose that the time period available for colonization of a host is an important determinant for successional symbiont colonization and persistence and is extremely relevant for symbiont transfer amongst hosts in varied developmental stages. In other words, the distribution of developmental stages of hosts in the host population could influence symbiont transfer and persistence. These findings are particularly interesting owing to their applicability to other similar symbiont metacommunities such as gut microbial symbionts. For example, in many mammals, it is well known that certain essential microbes are transferred into the alimentary tract of the offspring during lactation, thereby being acquired only during these early stages of growth and are then harbored for life (Gilbert, 2014). However, other microbial symbionts may be acquired during broader windows at various stages of host growth and through various diets or other sources (Mcfall-Ngai, 1994; Walter & Ley, 2011). Therefore, such differences in the colonization window lengths of symbionts could make certain symbionts more vulnerable to extinction than others with consequences for the persistence and structure of the symbiont metacommunities.

Our simulation results also showed that certain features of symbiont metacommunities are similar to non-symbiont metacommunities. That colonization success of predators never exceeded that of their best performing galler prey is in line with the trophic rank hypothesis which postulates that species diversity reduces while moving up the trophic level owing to additional requirements of prey availability for predators (Srivastava, Trzcinski, Richardson & Gilbert, 2008; Losos & Ricklefs, 2009). In all our simulations, total prey occurrence increased with increasing cumulative OW lengths of the prey. Therefore, the colonization success of predators correspondingly increased irrespective of actual diet breadth or the success and persistence of single galler prey. Though this particular result may have likely occurred because we did not capture predator–prey population dynamics, it still emphasises the inherent importance of prey occurrence rather than just diet breadth of predators for dictating predator colonization success. Specifically, intensity of predation on prey, and how reduced prey availability influences predator persistence in metacommunities need to be addressed.

Finally, in all our results, increased host abundances led to increased metacommunity persistence. This is in agreement with island biogeography theory; increasing numbers of microcosms/refugia/islands support increased species diversity and abundance (Losos & Ricklefs, 2009).

A few assumptions in our model require may require reconsideration for future work. All symbionts were assumed to have the same dispersal ability. However, our own experimental investigations of *F. racemosa* symbiont wasps suggest that predators of the non-mutualistic gallers (competitors) exhibit reduced dispersal capacities (Venkateswaran et al., 2017). We speculate that increased host abundances are required for the persistence of these predators (apparent mutualists) with reduced dispersal kernels (Herrera, Morales, & García, 2011) and therefore the minimum tree numbers for the persistence of these apparent mutualists that we deduce are likely underestimates. Incorporation of population dynamics within such communities is a next step in this investigation. Future work should also test the impact of dispersal limitation to better understand colonization success. Wasp species also vary in colonization pressures through colonization windows and trophic positions which can select for different dispersal abilities (Venkateswaran et al., 2017). We are far from understanding the actual distances that wasps traverse in space, or the time they have available for dispersal. We believe that an investigation of these factors is critical in order to best understand the scale at which to situate an empirical study. Generally, our investigation provides a novel framework and important insights for symbiont metacommunity membership in natural microcosms.

## Acknowledgements

We would like to thank Rohini Balakrishanan and Priya Iyer for providing valuable insights at various stages of the conceptualization of the idea and model development. We also thank Krishnapriya Tamma, Karpagam Chelliah, Kavita Isvaran, Yuvraj Ranganathan, Mahua Ghara and Anusha Krishnan for helpful input while drafting this paper. We thank Priyanka Ambavane for the artwork, and Sumitra Shankaran and Jaideep Joshi for providing assistance with code and simulations. The work was supported by Ministry of Human Resource Development—Indian Institute of Science for graduate scholarship.

## Author’s contributions

VV conceived the study, VV ran the simulations and analyzed the data, VV and RMB were involved in jointly writing the manuscript and reviewing it and the data for important intellectual content.

## Data accessibility

All simulated data and the phenology data will be deposited in a suitable online repository such as Dryad or any other suitable repository upon publication.

## Competing interests

The authors declare that they have no competing interests

## Funding

The project utilized funds from the Department of Science and Technology (DST), DST-FIST, Department of Biotechnology–Indian Institute of Science Partnership Support, and Ministry of Environment, Forests & Climate Change, Government of India.

## Ethical treatment of animals

all research presented in the manuscript was conducted in accordance with all applicable laws and rules set forth by their governments of India and the Indian Institute of Science.

## Supplementary

### Natural history and membership constraints in fig wasp communities

#### a) Inter-symbiont associations and dependencies

Galls are niches that are co-constructed by a host-plant and symbiont (typically an insect); the process of gall formation is referred to as galling, and the gall inducer a galler. Each fig wasp community contains a mutualist that both galls and pollinates the thousands of uniovulate flowers within globular, urn-shaped and enclosed inflorescences called syconia (Janzen, 1979). Galling occurs during egg deposition into floral tissue which then induces gall formation; offspring are sheltered and nourished within these galls. Typically, only a single offspring develops within a gall, and galling spaces within a single fig inflorescence are limited. Other non-pollinating gallers compete with the pollinator for galling space in floral tissues but offer no pollination service to the plant (Janzen, 1979; Machado, Robbins, Gilbert, & Herre, 2005). Parasitoids arrive a few days after galler oviposition and oviposit in those galls containing galler offspring. Parasitoid offspring subsequently feed on the developing prey within galls (Janzen, 1979; Borges, 2015). Parasitoids in fig wasp communities require the appropriate host species developing within the fig syconium at the appropriate developmental stage (Yadav & Borges, 2017). Therefore, the availability of appropriate host-derived habitats for gallers and the availability of gallers for parasitoids are likely important membership constraints for these communities.

#### b) Symbionts in mutualism function and persistence

The persistence of the mutualism between the fig and the pollinating wasp is central; since the relationship between the mutualist and host is bi-directional and obligate, the breakdown of this relationship could lead to community collapse (Bronstein, Gouyon, Gliddon, Kjellberg, & Michaloud, 1990). This can happen in one of two ways in fig systems: a) failure to pollinate syconia that often leads to their abortion (Anstett, Kjellberg, & Bronstein, 1996) which should make these ‘host-derived’ habitats unavailable for the entire wasp community, and b) over time, pollinator extinction should result in the eventual extinction of the host plant, which would also lead to community collapse (Bronstein, Gouyon, Gliddon, Kjellberg, & Michaloud, 1990; Anstett, Michaloud, & Kjellberg, 1995; Harrison & Yamamura, 2003). A few species in the community are detrimental for the mutualism. For instance, the galler species that compete with the pollinator are detrimental as they are expected to compete for space with the pollinator and do not offer pollination services (Krishnan et al., 2015). Certain parasitoid species that specialize and feed on the pollinator offspring may also be detrimental to the mutualism (Krishnan et al., 2015; Yadav & Borges, 2017). However, certain other parasitoid species specialize only on competing gallers, reducing the competition for the mutualist and thereby indirectly benefitting the mutualism (apparent mutualists) (Krishnan & Borges, 2014; Krishnan et al., 2015). For a complete representation of host–community interactions and dependencies refer to Figure S1.

#### c) Phenology and resource dispersion for symbionts

The fruiting phenologies of figs dictate habitat availability for different fig wasp species within communities. Tropical fig species have aseasonal fruiting phenologies (Anstett et al., 1995; Krishnan, Pramanik, Revadi, Venkateswaran, & Borges, 2014). The phenology of each syconium passes through a series of developmental phases followed by a stage of quiescence called the gap phase (Figure 2). Individual fig trees often exhibit reproductive synchrony; all syconia within a tree are usually in the same developmental phase (Bronstein et al., 1990). However, in a given population, trees are often reproductively asynchronous with each other. These features create unique resource initiations in space; different trees are usually in different phenological stages at a given time (Bronstein et al., 1990). Each wasp oviposits in the fig inflorescence during a specific window during the development of the fig (also called the oviposition window (OW)). The length and position of this window is deduced from natural observations of the timing of wasp arrival and is observed to be unique for each wasp species; the length appears to be dictated by how long resources last in the syconium for the wasp species in consideration (Ghara & Borges, 2010; Yadav & Borges, 2017). These temporal phenological features of the syconium and the tree not only a) ensure that the natal trees bears no suitable syconia that can be colonized by the wasps emerging from them (a feature that forces wasps to leave their natal trees), but also b) dictate the patchiness of the oviposition resources for each member of the fig wasp community (Bronstein et al., 1990; Venkateswaran, Shrivastava, Kumble, & Borges, 2017). In this way, both the habitat availability for each wasp species and wasp release are intricately linked with the ontogeny of the syconium (the fruiting phenology of the tree). Therefore, differences in resource dispersion or dispersal limitation can be viewed as another constraint for membership.

### Estimating lengths of phenological phases from field data

The data set used consisted of approximately 20 bunches per tree (n = 17 trees) that were marked and observed over multiple fruiting cycles (∼10 fruiting cycles). Observations were conducted every alternate day for 643 days (∼2 years). Doubly-flanked phases, where possible, were identified to conservatively estimate phase lengths. We defined complete phases as those phases that were flanked on both ends by phases that precede or follow the phase in consideration. Since no phase preceded the pre-receptive stage and all syconia were recorded from the first day of initiation, we used only those pre-receptive stages that were followed by a receptive stage to determine pre-receptive stage length. To estimate the length of the gap phase, we calculated the difference between the end of fruit-ripening of one cycle and the beginning of the pre-receptive stage of the next cycle. Using the statistical program R (3.3.1), the length of each phase from this dataset was extracted. Instances of natural syconia abortion or failure due to other unaccounted reasons were not used for estimation of phase lengths (Figure S2).

### Order of arrival of galler species

The temporal ordering of OW only influenced the number of trees harboring that particular wasp species at any time (Figure S5 a–j); this was especially evident at tree abundances greater than 50. As the number of trees were increased to support all wasp species stably, the earlier the OW, the more number of trees on any day that incubated the wasp species in the community (Figure S5 e–j). However, since both PCS and CO are not different in Model 1, the number of trees holding a wasp species does not seem to confer it any advantage in terms of colonization success; earlier colonizing wasps have longer periods of association with the host.

### Sensitivity analysis

The sensitivities of our models were tested to ensure that OW lengths positively impacted colonization success of gallers and the performance of parasitoids specializing on that galler. A single galler and a single parasitoid species were used in these simulations. Galler OWs were changed as a fraction of the pre-receptive phase (from the first 1/10th to the entire pre-receptive phase, in increments of 1/10ths). The parasitoid had a fixed oviposition window (5/10th to 8/10th of the developmental phase). The performance of both the galler and parasitoid were assessed by running 30 iterations for every value of OW change for the galler.

The sensitivity analysis revealed that with systematically increasing oviposition window lengths, gallers were better colonizers in the simulations as expected (Figure S6). Parasitoid colonization depended on the colonization success of the galler as expected. Specifically, consistent with the previous models described, oviposition length positively influenced PCS values; increasing galler oviposition windows did not affect parasitoid PCS (Figure S6a).

Oviposition window lengths also positively influenced CO for gallers which increased the CO for parasitoids and the PCS of the latter were never more than that of the former indicating galler colonization success limited parasitoid colonization success through CO (Figure S6 b). Finally, the sensitivity analyses revealed that our model was sensitive to the length of the oviposition window, a parameter of importance for the investigation.

### Pseudocode

The pseudocode presented below represents an overview of the main code and is intended as a guide for the main code in Appendix-2 (R script). We suggest referring to code in the R studio console. Information on the structure of the master sequence and the data frame and their roles are also provided. The first code provided documents the code used to investigate the *F. racemosa* wasp community performance (the real community). All other model variants can be performed by varying the relevant parameter values with this same simulation framework. In cases were the comparisons were made between theoretical gallers or parasitoids (the model variants in the main manuscript), pollinators were also present to ensure pollination requirement for resource persistence.

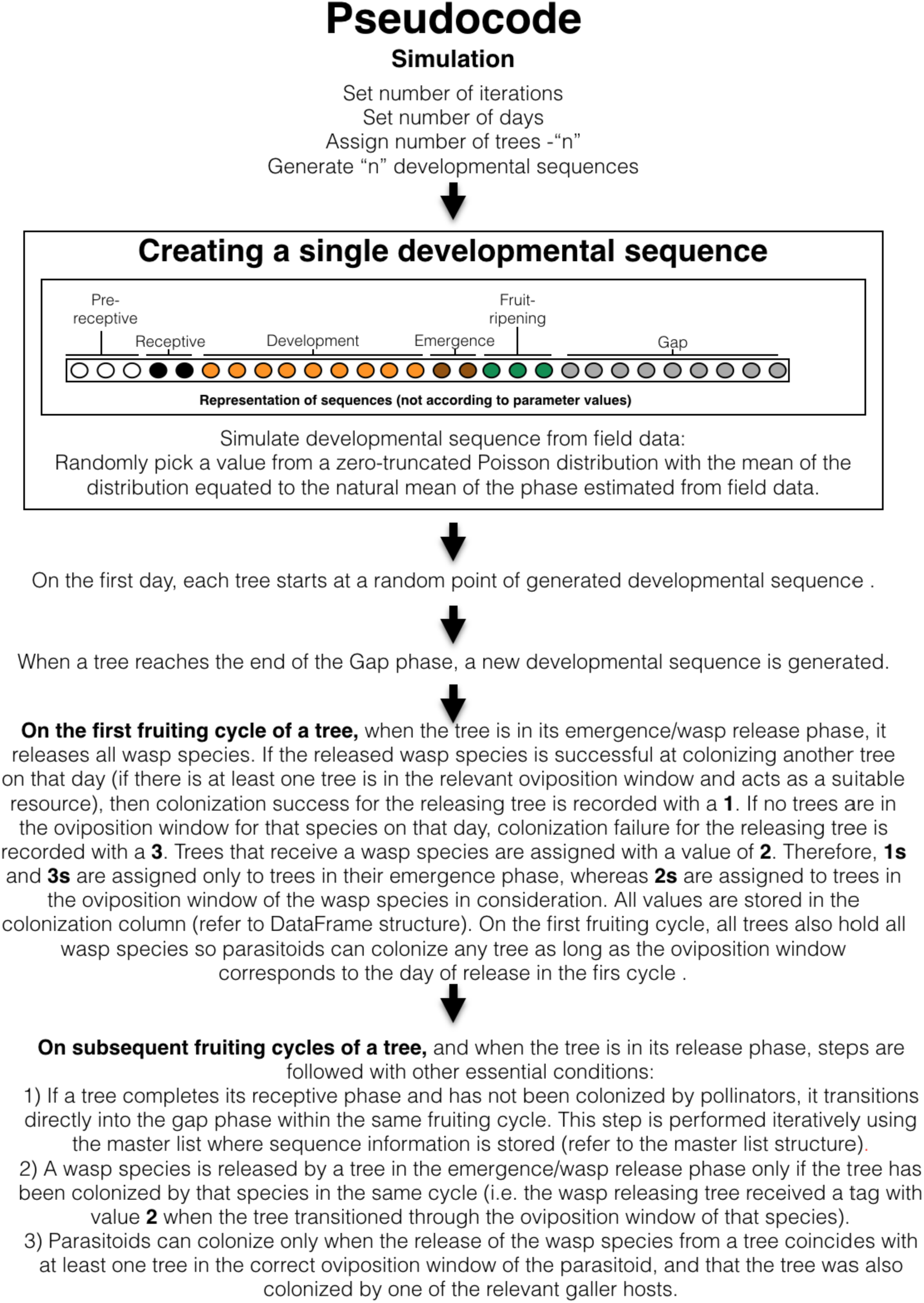
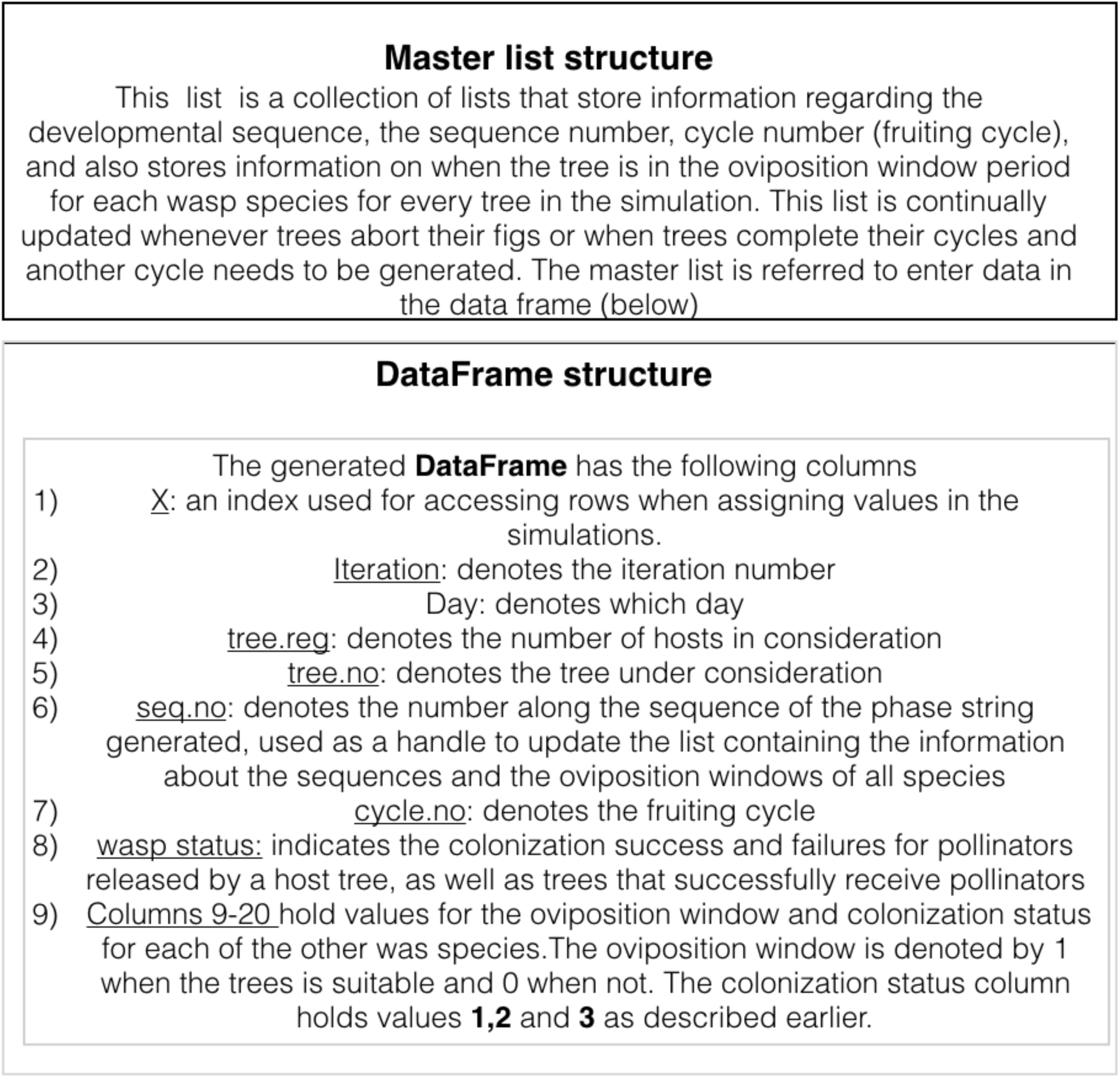

---

**Figure S1:**
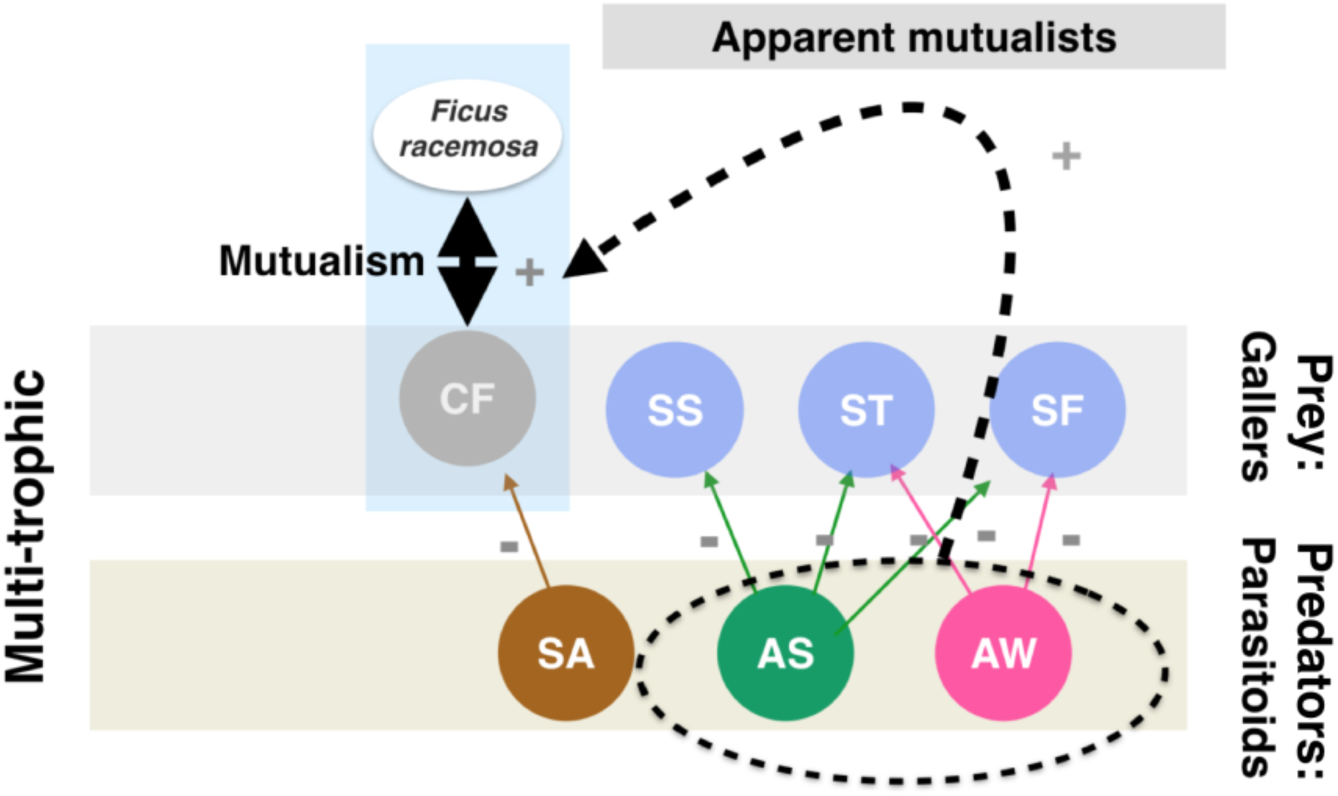
The trophic network of the wasp community of *Ficus racemosa*. Species abbreviations are AS-*Apocrypta* species 2, AW-*Apocrypta westwoodi*, SS-*Sycophaga stratheni*, ST-*Sycophaga testacea*, SF-*Sycophaga fusca*, SA-*Sycophaga agraensis*, CF-*Ceratosolen fusciceps*.

**Figure S2:**
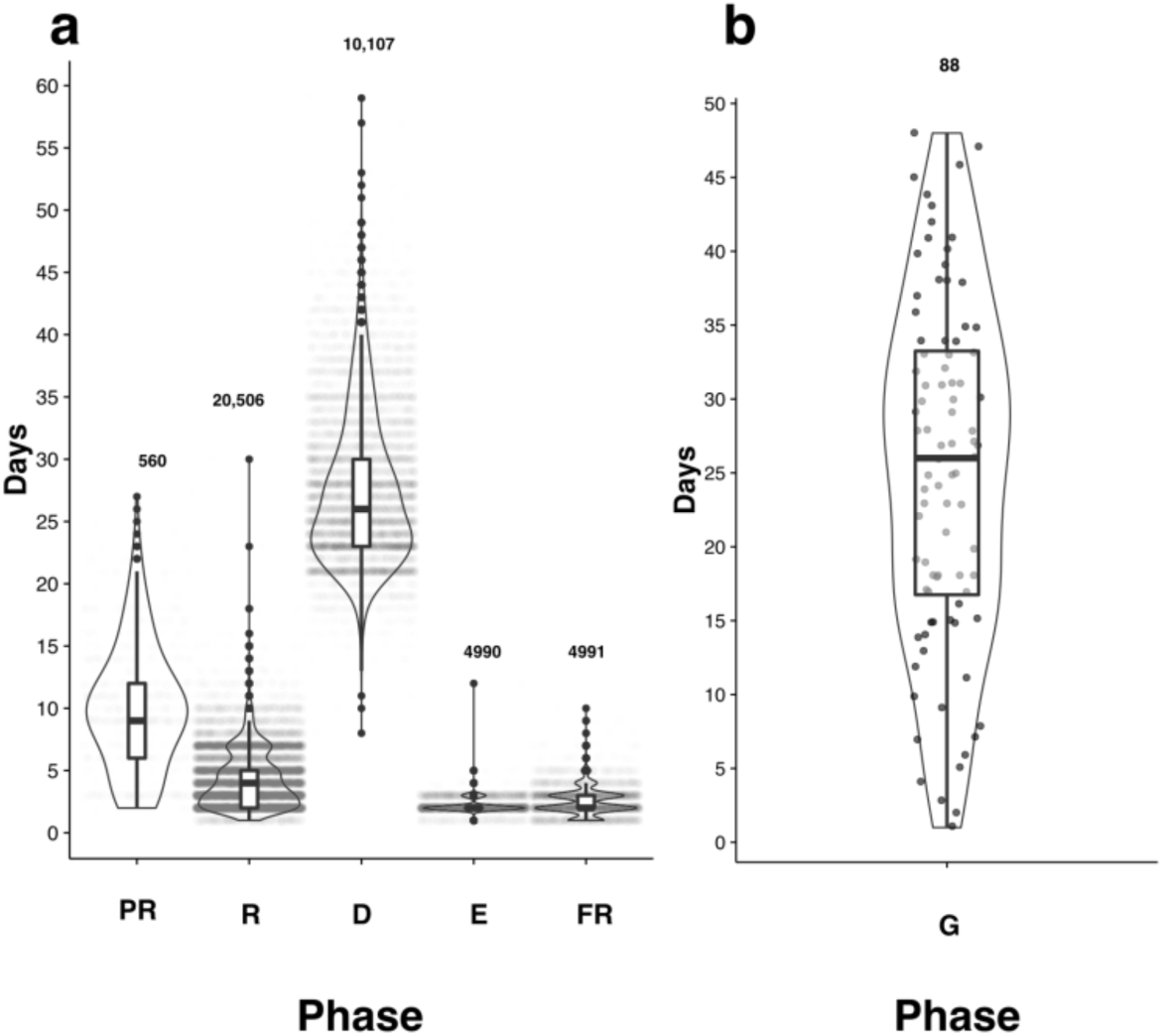
The length of each phenological phase of *F. racemosa* syconia a) Depicts the spread of each developmental phase of the syconium. Phase abbreviations are PR: pre-receptive; R: receptive; D: development; E: wasp emergence; FR: fruit ripening. Numbers indicate sample size. Greater value overlaps are represented by darker colors. Within the violin plots are box plots. Dark circles indicate the outliers. b) Depicts the spread associated with the G phase or the quiescent phase between two fruiting cycles; G was estimated differently (see methods in Appendix 1).

**Figure S3:**
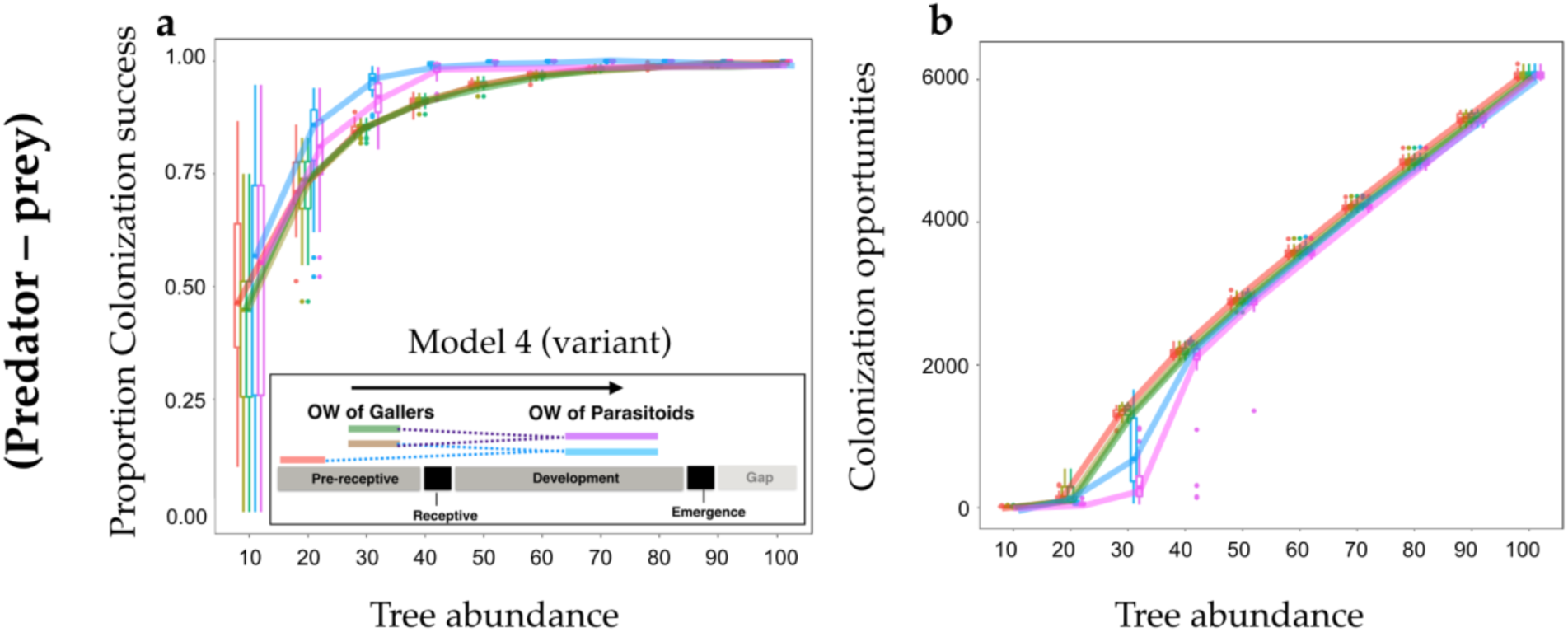
A variant of Model 4; performance of parasitoids (predators) that are dependent on the same number of prey species, but that differ in the staggering of oviposition windows of their prey. Details of parameters values of this model variant indicated in Table S5.

**Figure S4:**
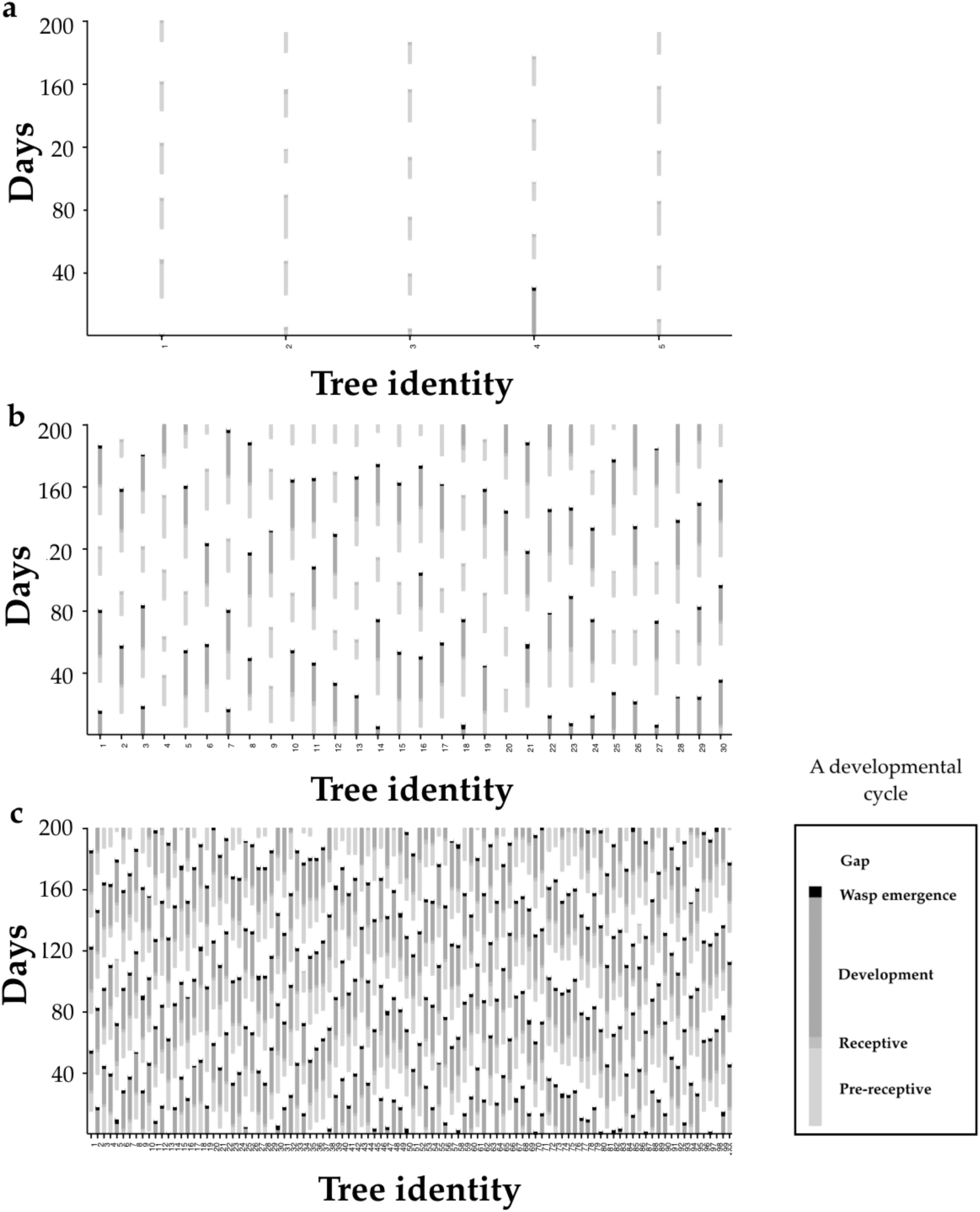
The developmental trajectories of syconial microcosms on individual fig trees in different tree regimes over 200 days. Trees abort figs when pollinators fail to colonize. In a) with only 5 trees, the frequency of resource presentation and persistence is zero. In b) with 30 trees, and in c) with 100 trees, the number of abortions were reduced.

**Figure S5:**
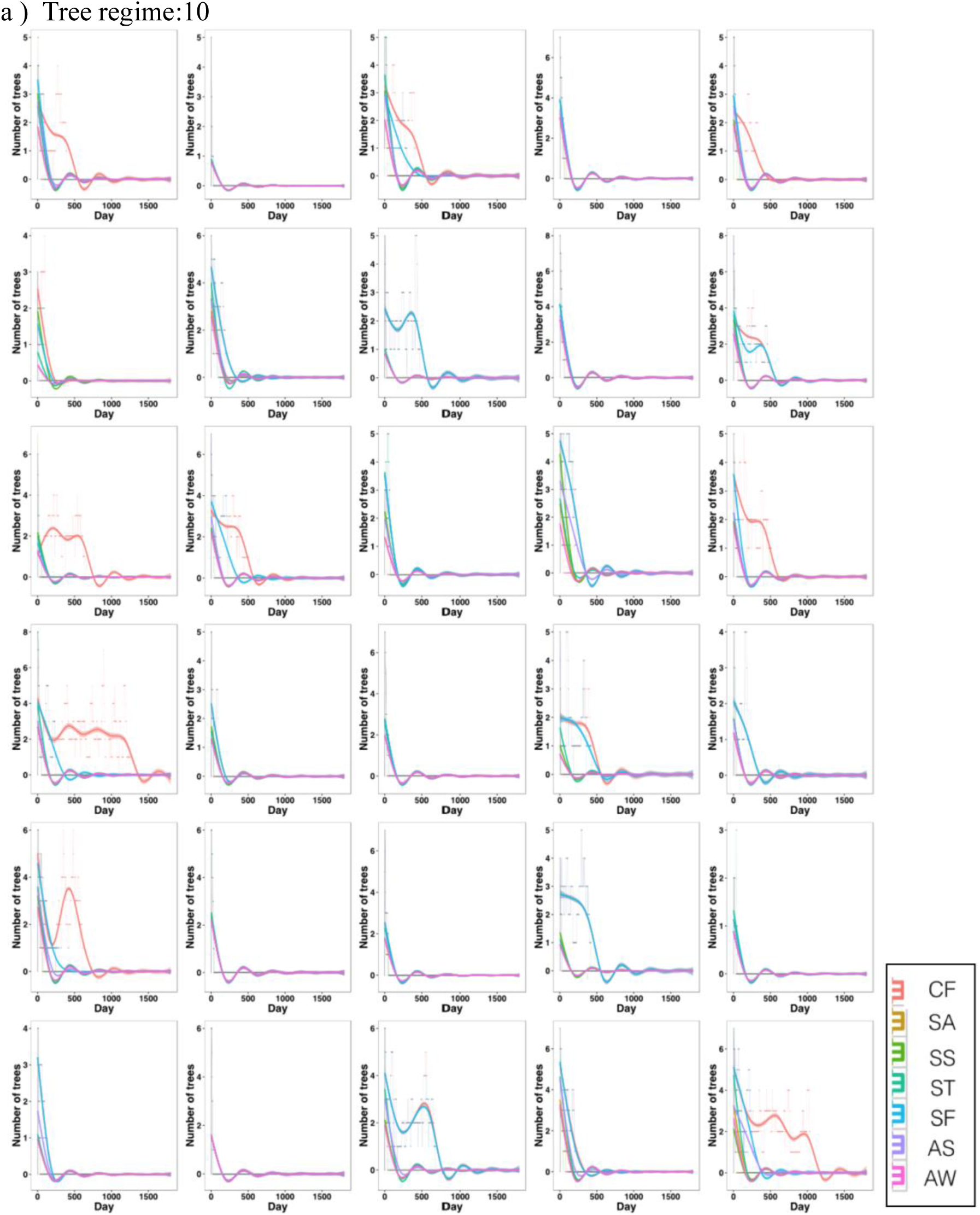
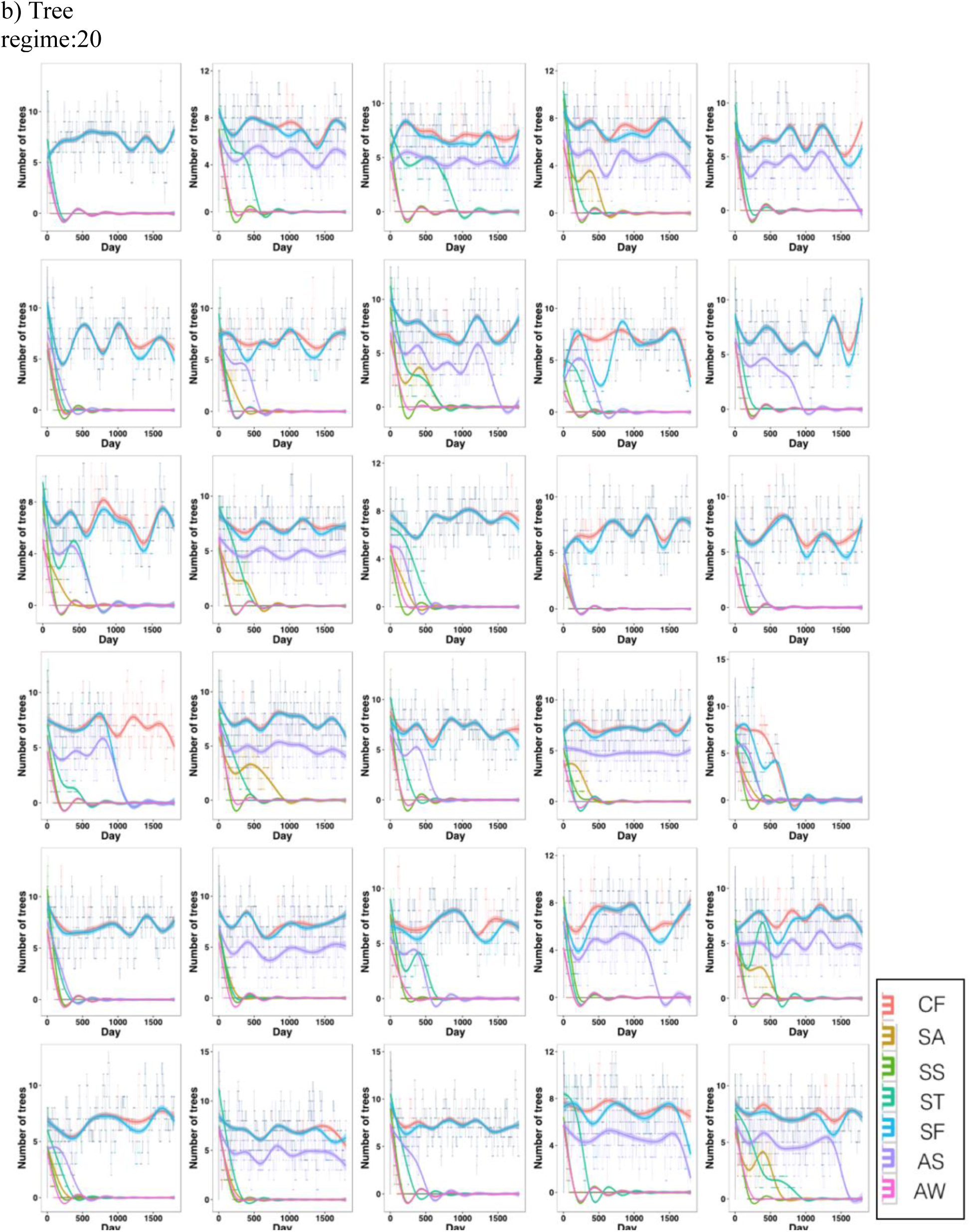
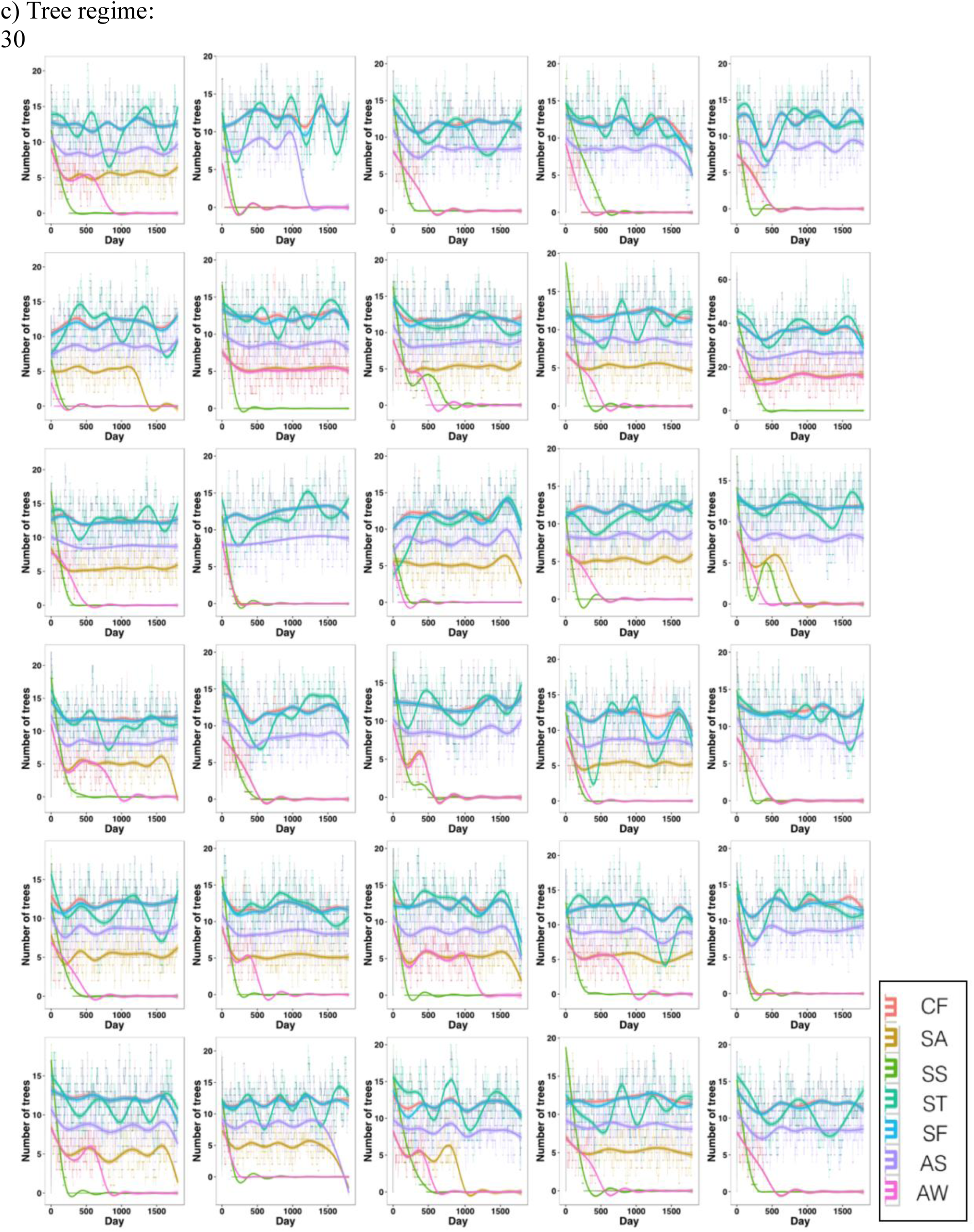
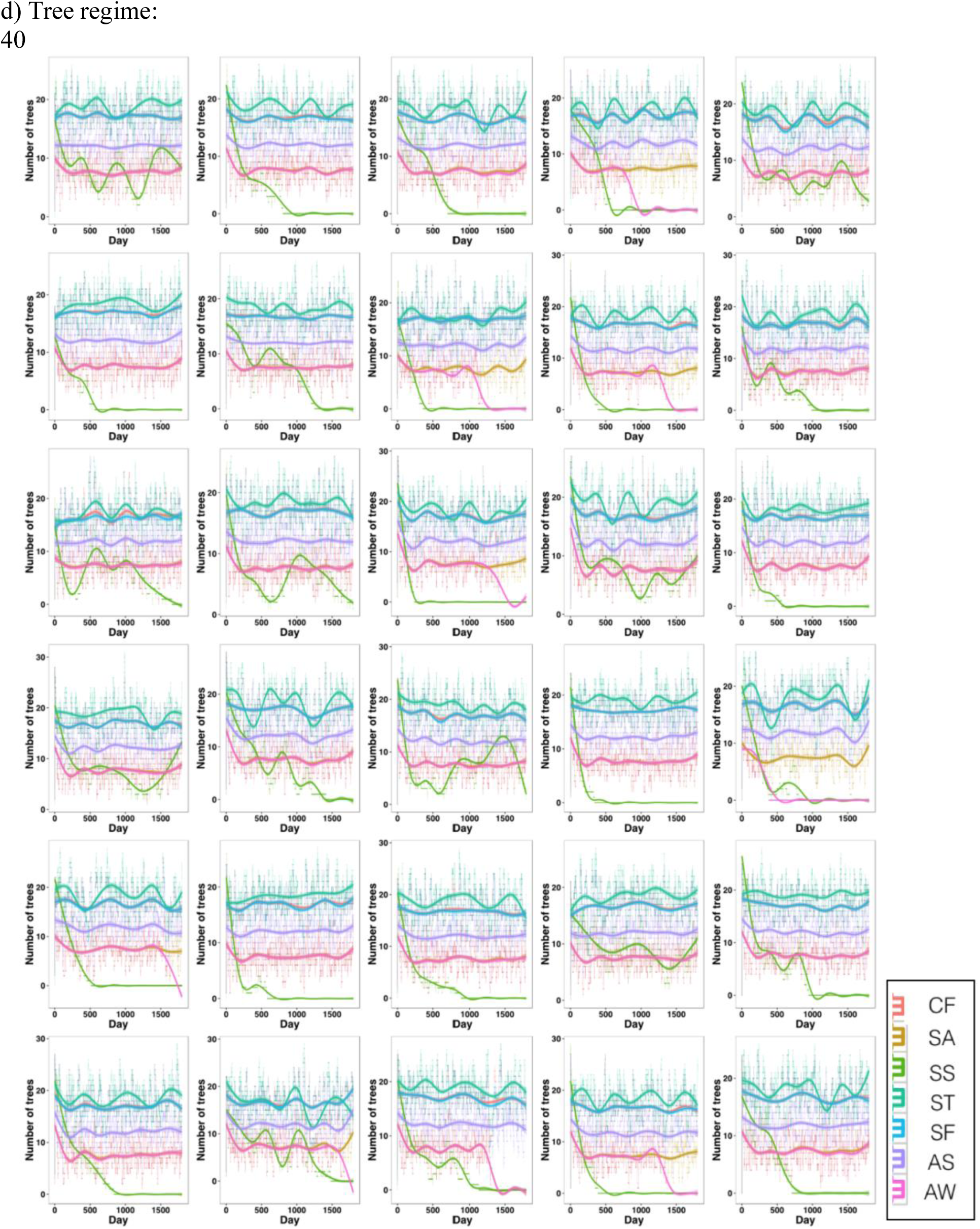
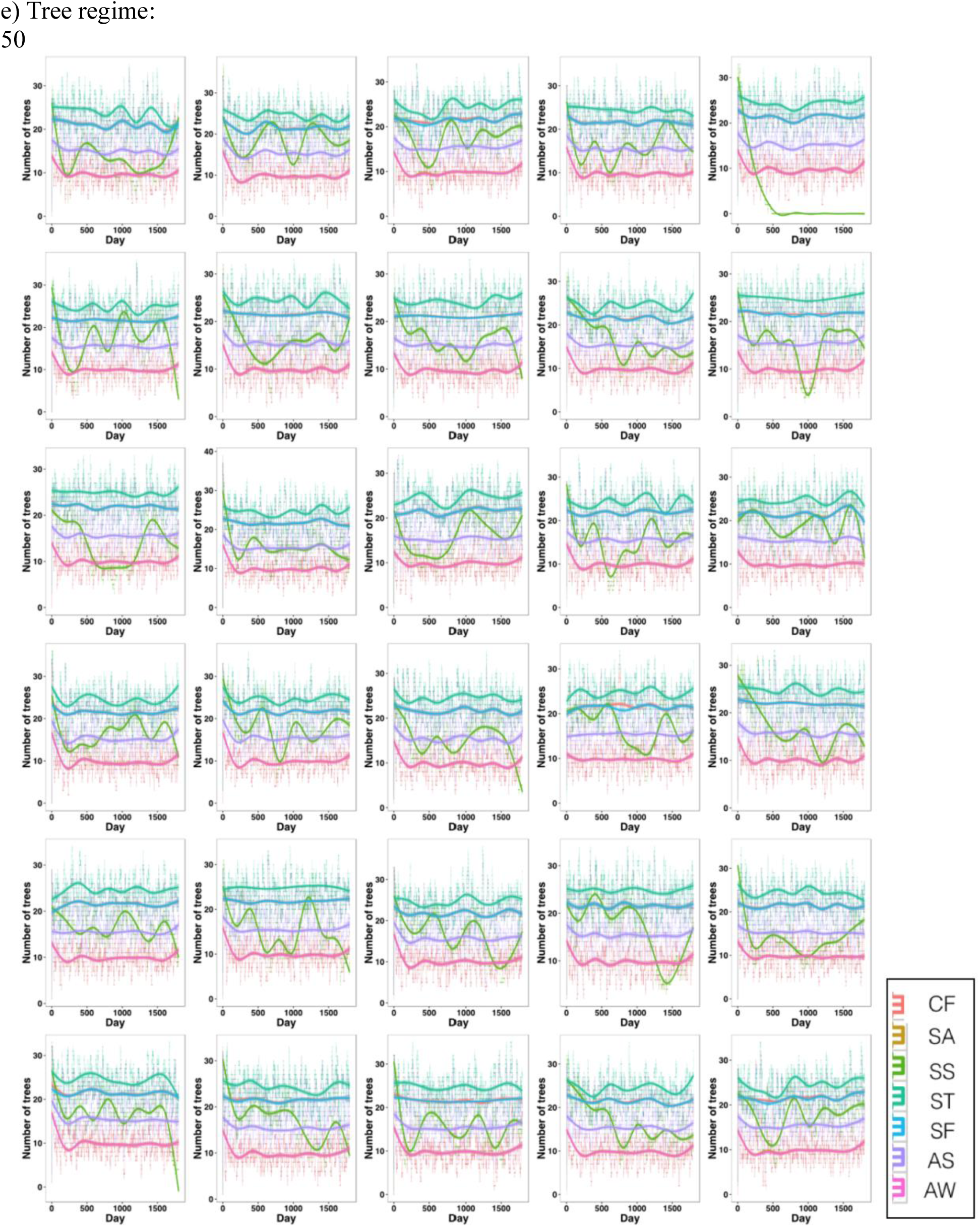
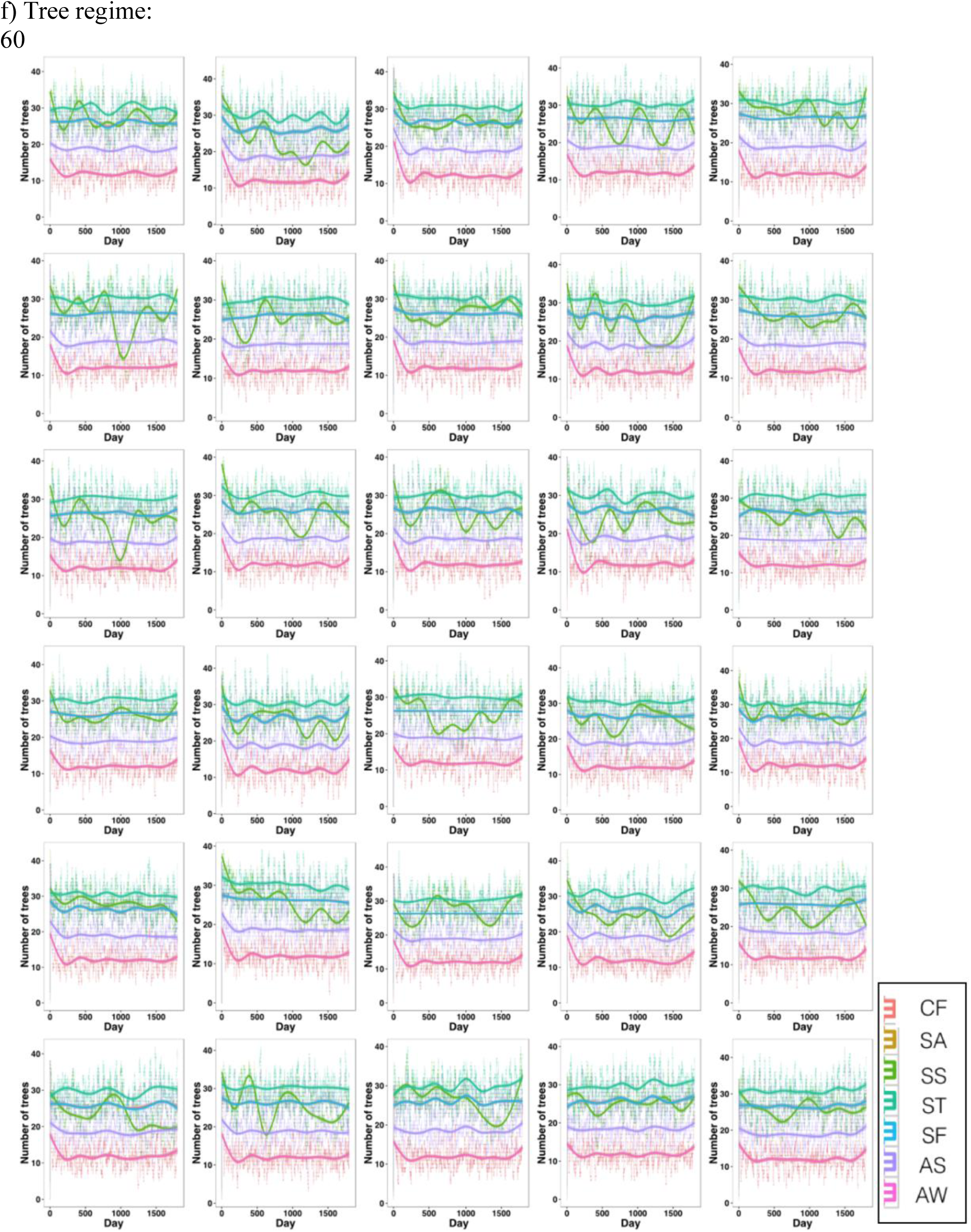
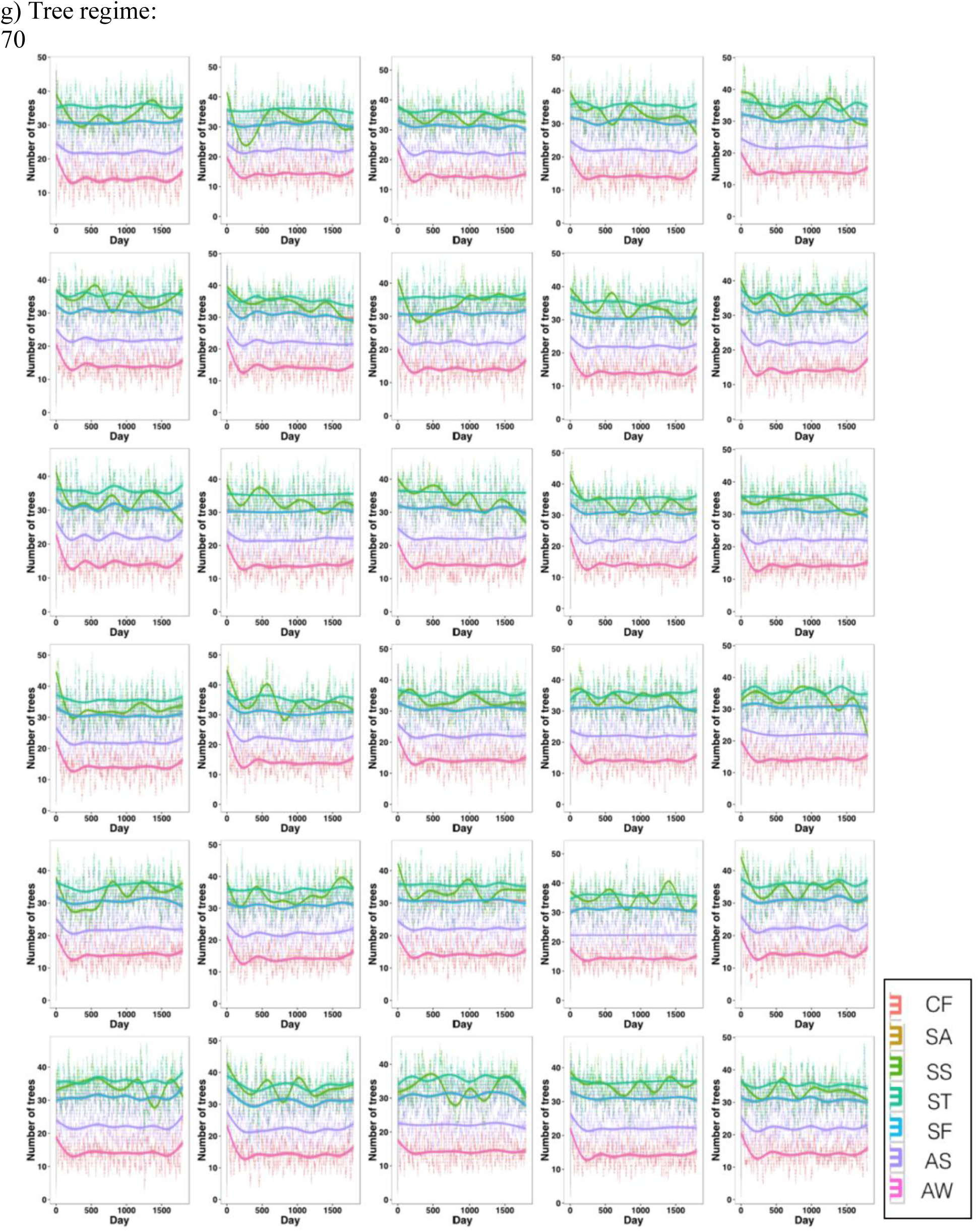
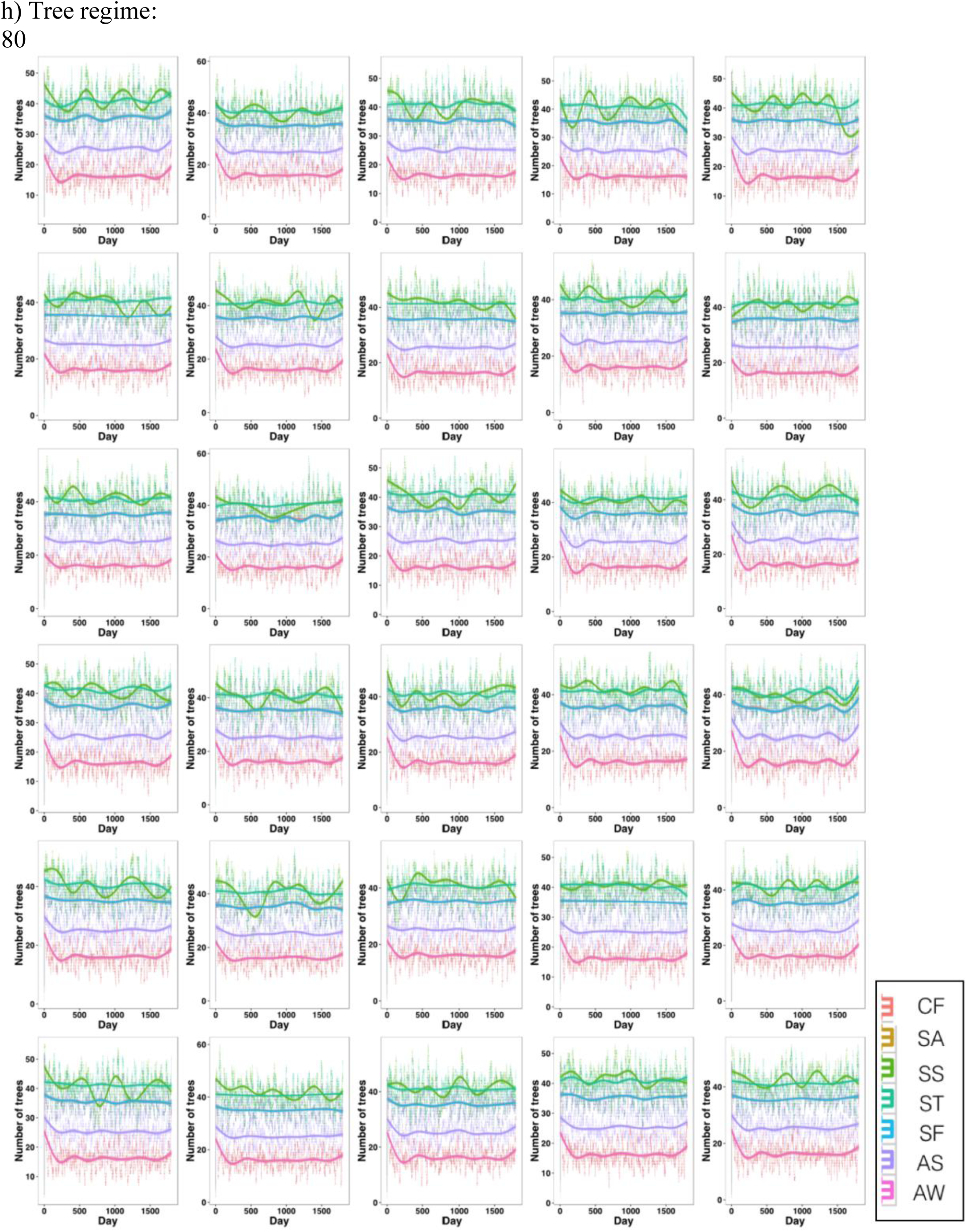
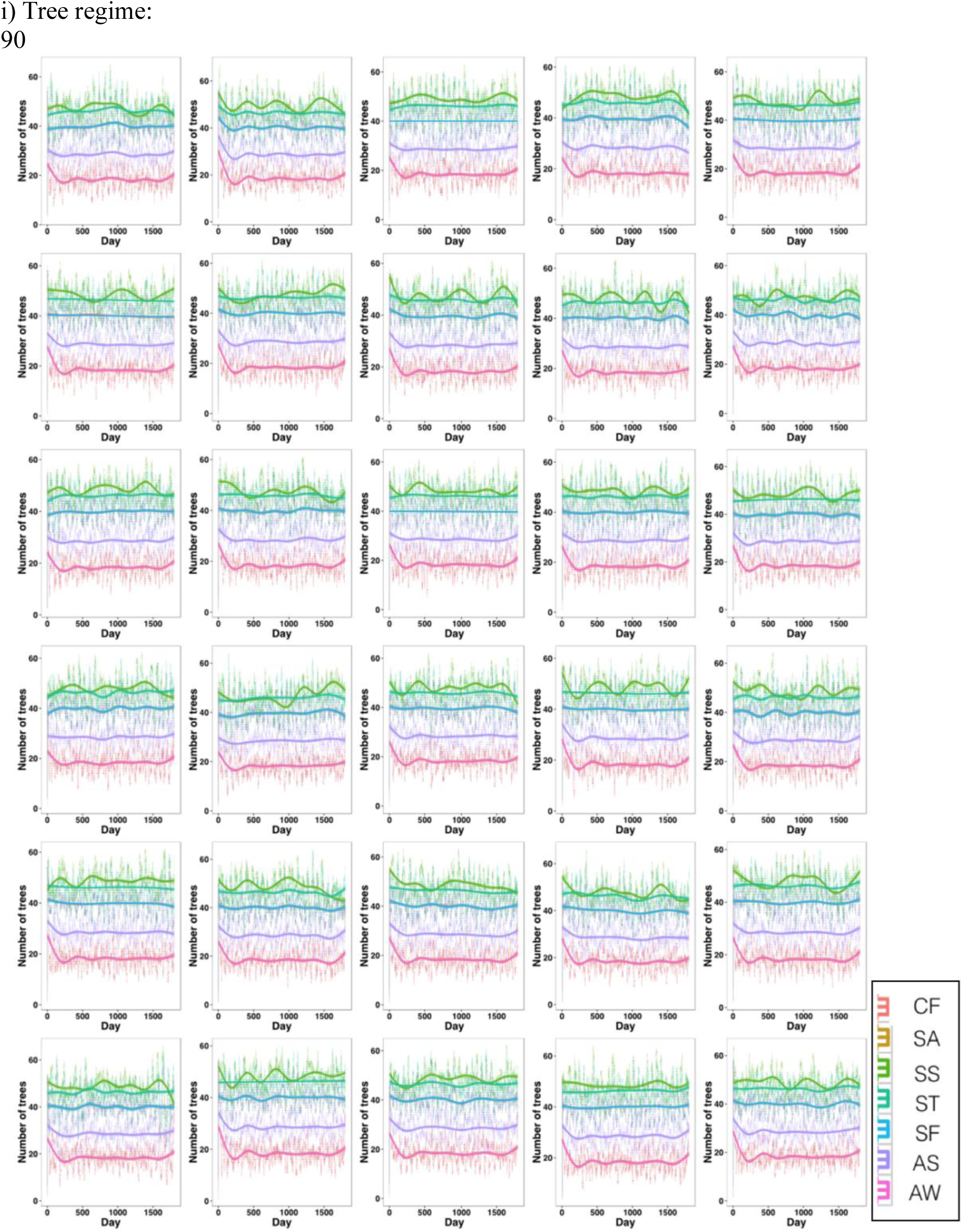
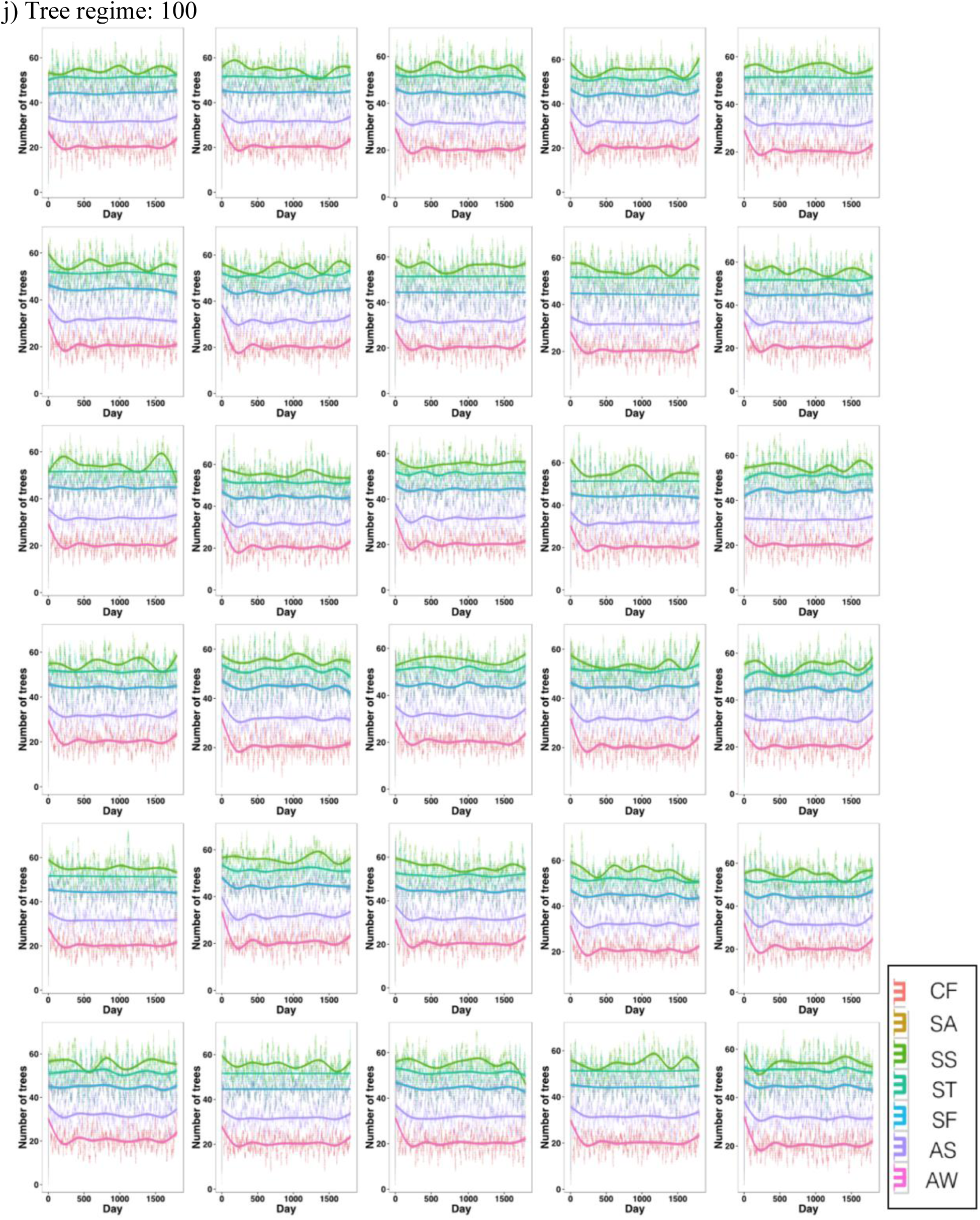
The number of trees in each simulation harboring each wasp species over 1800 days. The species abbreviations are the same as in Fig S1). Each panel (a-j) represents one tree abundance regime and depicts each of the 30 iterations. Loess lines are plotted to aid visualization. a) Tree regime:10

**Figure S6:**
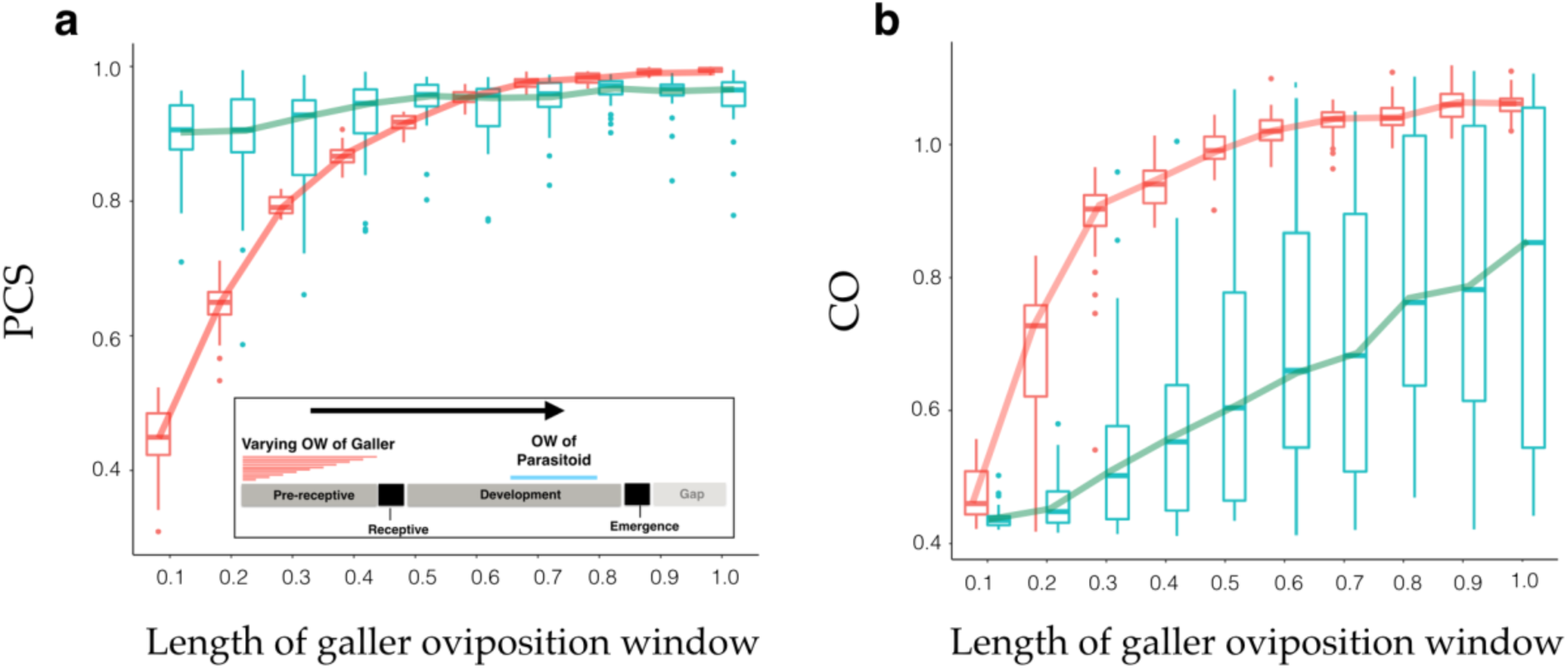
Sensitivity analysis to assess how the length of the oviposition window influences galler performance (red) and the performance of a parasitoid (green) that feeds on the galler. Only the oviposition length of the galler is varied while that of the parasitoid is kept constant. The length of the oviposition window for the galler in the x-axis indicates the fraction of length of the simulated pre-receptive phase (see above for details).

**Table S1:** Summary statistics associated with each phenological phase of *F. racemosa* in days.

| Phase | Mean | Variance | Range |
| --- | --- | --- | --- |
| Pre-receptive | 9.975 | 23.78 | (2-27) |
| Receptive | 4.238418 | 4.85 | (1-30) |
| Development | 27.33205 | 32.71 | (8-59) |
| Emergence | 2.15434 | 0.25 | (1-12) |
| Fruit ripening | 2.495138 | 1.11 | (1-10) |
| Gap | 25.056 | 136.79 | (1-48) |

**Table S2.** Summary of the results of Models 1 to 4.

| **Model**<br>**(Predictor)** | **Response**<br>**variable** | **Difference in**<br>**Proportion**<br>**Colonization**<br>**Success**<br>**(PCS)** | **Tree**<br>**regimes**<br>**with**<br>**statistically**<br>**significant**<br>**differences** | **Difference in**<br>**Colonization**<br>**Opportunities**<br>**(CO)** | **Tree**<br>**regimes**<br>**with**<br>**statistically**<br>**significant**<br>**differences** |
| --- | --- | --- | --- | --- | --- |
| Model 1<br>(Position of<br>OW) | Galler<br>colonization | No | Not<br>applicable | No | Not<br>applicable |
| Model 2<br>(Length of<br>OW) | Galler<br>colonization | Yes | 10–100 | Yes | 20–100 |
| Model 3<br>(Degree of<br>parasitoid<br>specialization) | Parasitoid<br>(predator)<br>colonization | No | Not<br>applicable | No | Not<br>applicable |
| Model 4<br>(Differences in<br>galler<br>performance) | Parasitoid<br>(predator)<br>colonization | No | Not<br>applicable | Yes | 30–70 |

OW = oviposition window

**Table S3a:** Wilcoxon values and corresponding P values for Model 1 and Model 2 for comparing the PCS of the galler species

| Number of trees | Model 1 |  | Model 2 |  |
| --- | --- | --- | --- | --- |
|  | W | P | W | P |
| 10 | 433.5 | 0.8129 | 144.5 | < 0.0001 * |
| 20 | 355 | 0.1635 | 16.5 | < 0.0001 * |
| 30 | 143 | < 0.0001 * | 0 | < 0.0001 * |
| 40 | 207 | 0.0002 | 0 | < 0.0001 * |
| 50 | 183 | < 0.0001 * | 0 | < 0.0001 * |
| 60 | 215 | 0.0004 | 0 | < 0.0001 * |
| 70 | 255 | 0.0035 | 0 | < 0.0001 * |
| 80 | 227.5 | 0.0010 | 0 | < 0.0001 * |
| 90 | 273 | 0.0084 | 0 | < 0.0001 * |
| 100 | 160 | < 0.0001 * | 0 | < 0.0001 * |

**Table S3b:** Wilcoxon values and corresponding P values for Model 1 and Model 2 for comparing the CO for each species (gallers) at each tree regime

| Number of trees | Model 1 |  | Model 2 |  |
| --- | --- | --- | --- | --- |
|  | W | P | W | P |
| 10 | 431 | 0.7839 | 364.5 | 0.2079 |
| 20 | 383.5 | 0.3291 | 173 | < 0.0001 * |
| 30 | 310 | 0.0392 | 9.5 | < 0.0001 * |
| 40 | 282.5 | 0.0135 | 0 | < 0.0001 * |
| 50 | 359 | 0.1809 | 0 | < 0.0001 * |
| 60 | 336 | 0.0933 | 0 | < 0.0001 * |
| 70 | 413 | 0.5894 | 0 | < 0.0001 * |
| 80 | 407 | 0.5297 | 0 | < 0.0001 * |
| 90 | 433 | 0.8073 | 5 | < 0.0001 * |
| 100 | 426.5 | 0.7337 | 102.5 | < 0.0001 * |

**Table S3c:**
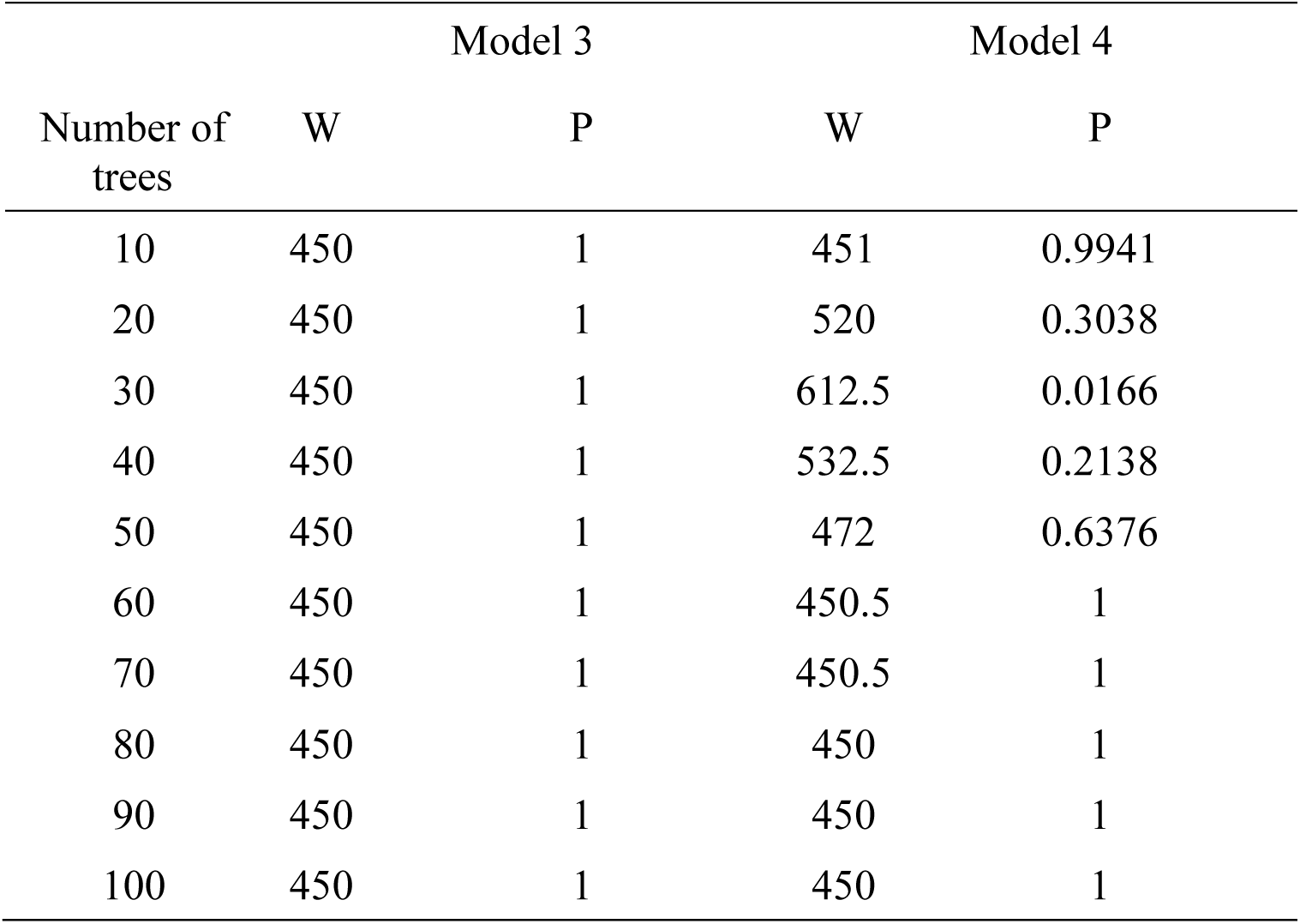
Wilcoxon values and corresponding P values for Model 3 and Model 4 for comparing the PCS for each of the two species (parasitoids) at each tree regime

**Table S3d:** Wilcoxon values and corresponding P values for CO comparisons for each of the two species (parasitoids) at each tree regime for Model 3 and Model 4.

| Number of trees | Model 3 |  | Model 4 |  |
| --- | --- | --- | --- | --- |
|  | W | P | W | P |
| 10 | 450 | 1 | 459 | 0.8997 |
| 20 | 450 | 1 | 512.5 | 0.3589 |
| 30 | 450 | 1 | 673.5 | < 0.05 * |
| 40 | 450 | 1 | 633.5 | < 0.05 * |
| 50 | 450 | 1 | 566 | 0.0876 |
| 60 | 450 | 1 | 514 | 0.3477 |
| 70 | 450 | 1 | 487.5 | 0.5841 |
| 80 | 450 | 1 | 473.5 | 0.7337 |
| 90 | 450 | 1 | 460 | 0.8882 |
| 100 | 450 | 1 | 454 | 0.9587 |

**Table S4a:**
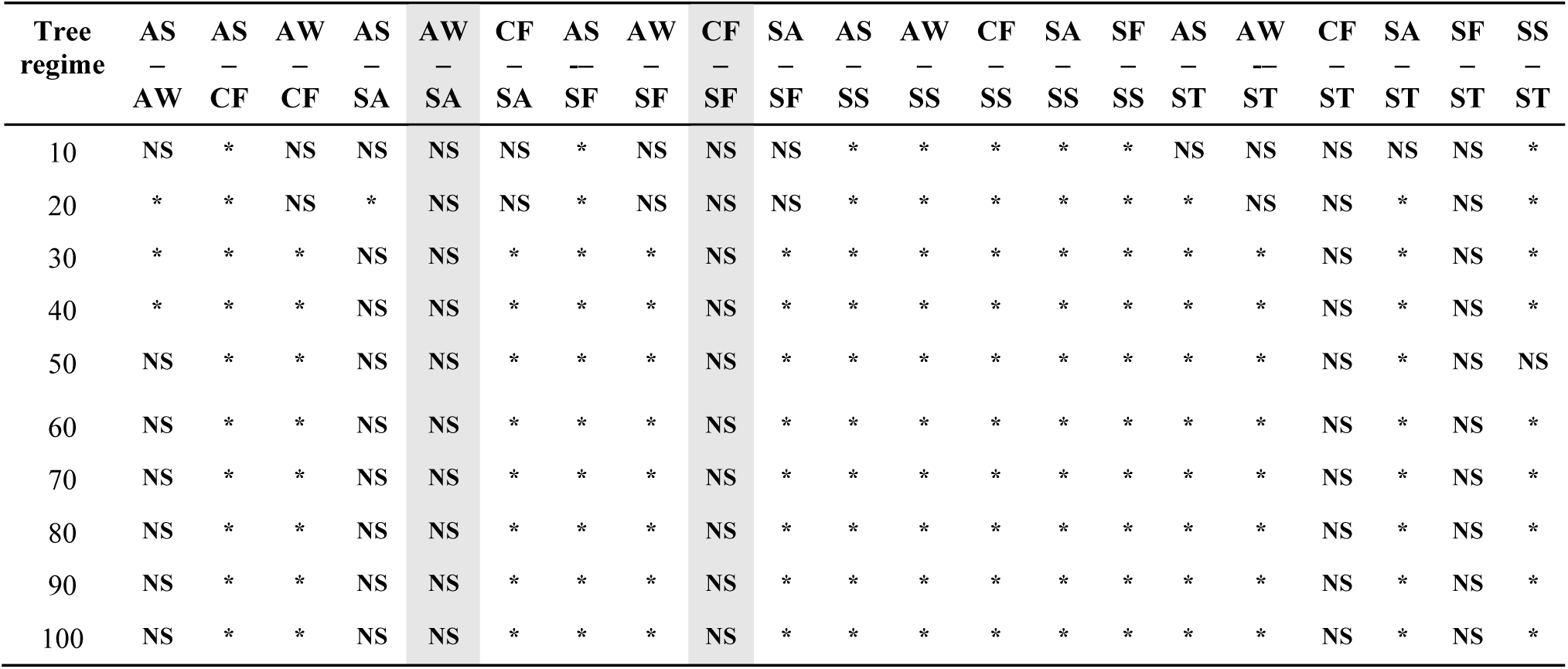
Dunn test comparisons with Bonferroni corrections (alpha=0.05) of PCS at different tree regimes. Shaded areas depict comparisons with consistently no difference at all tree regimes. Species abbreviations are as indicated in figure 1.

| Tree regime | AS<br>–<br>AW | AS<br>–<br>CF | AW<br>–<br>CF | AS<br>–<br>SA | AW<br>–<br>SA | CF<br>–<br>SA | AS<br>–<br>SF | AW<br>–<br>SF | CF<br>–<br>SF | SA<br>–<br>SF | AS<br>–<br>SS | AW<br>–<br>SS | CF<br>–<br>SS | SA<br>–<br>SS | SF<br>–<br>SS | AS<br>–<br>ST | AW<br>–<br>ST | CF<br>–<br>ST | SA<br>–<br>ST | SF<br>–<br>ST | SS<br>–<br>ST |
| --- | --- | --- | --- | --- | --- | --- | --- | --- | --- | --- | --- | --- | --- | --- | --- | --- | --- | --- | --- | --- | --- |
| 10 | NS | * | NS | NS | NS | NS | * | NS | NS | NS | * | * | * | * | * | NS | NS | NS | NS | NS | * |
| 20 | * | * | NS | * | NS | NS | * | NS | NS | NS | * | * | * | * | * | * | NS | NS | * | NS | * |
| 30 | * | * | * | NS | NS | * | * | * | NS | * | * | * | * | * | * | * | * | NS | * | NS | * |
| 40 | * | * | * | NS | NS | * | * | * | NS | * | * | * | * | * | * | * | * | NS | * | NS | * |
| 50 | NS | * | * | NS | NS | * | * | * | NS | * | * | * | * | * | * | * | * | NS | * | NS | NS |
| 60 | NS | * | * | NS | NS | * | * | * | NS | * | * | * | * | * | * | * | * | NS | * | NS | * |
| 70 | NS | * | * | NS | NS | * | * | * | NS | * | * | * | * | * | * | * | * | NS | * | NS | * |
| 80 | NS | * | * | NS | NS | * | * | * | NS | * | * | * | * | * | * | * | * | NS | * | NS | * |
| 90 | NS | * | * | NS | NS | * | * | * | NS | * | * | * | * | * | * | * | * | NS | * | NS | * |
| 100 | NS | * | * | NS | NS | * | * | * | NS | * | * | * | * | * | * | * | * | NS | * | NS | * |

**Table S4b:**
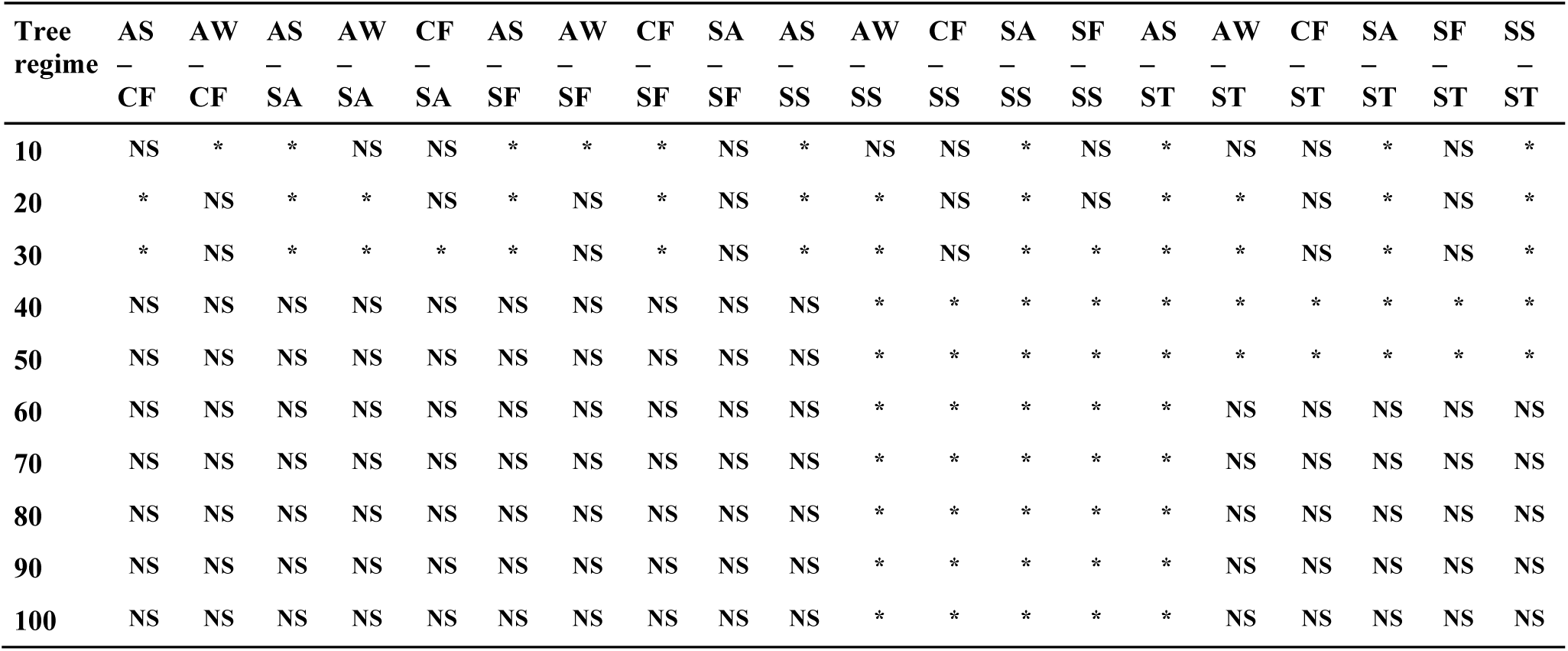
Dunn test comparisons of CO for all F. racemosa wasp species with Bonferroni corrections (alpha=0.05) at different tree regimes. Species abbreviations are as indicated in figure 1.

| Tree regime | AS<br>–<br>CF | AW<br>–<br>CF | AS<br>–<br>SA | AW<br>–<br>SA | CF<br>–<br>SA | AS<br>–<br>SF | AW<br>–<br>SF | CF<br>–<br>SF | SA<br>–<br>SF | AS<br>–<br>SS | AW<br>–<br>SS | CF<br>–<br>SS | SA<br>–<br>SS | SF<br>–<br>SS | AS<br>–<br>ST | AW<br>–<br>ST | CF<br>–<br>ST | SA<br>–<br>ST | SF<br>–<br>ST | SS<br>–<br>ST |
| --- | --- | --- | --- | --- | --- | --- | --- | --- | --- | --- | --- | --- | --- | --- | --- | --- | --- | --- | --- | --- |
| 10 | NS | * | * | NS | NS | * | * | * | NS | * | NS | NS | * | NS | * | NS | NS | * | NS | * |
| 20 | * | NS | * | * | NS | * | NS | * | NS | * | * | NS | * | NS | * | * | NS | * | NS | * |
| 30 | * | NS | * | * | * | * | NS | * | NS | * | * | NS | * | * | * | * | NS | * | NS | * |
| 40 | NS | NS | NS | NS | NS | NS | NS | NS | NS | NS | * | * | * | * | * | * | * | * | * | * |
| 50 | NS | NS | NS | NS | NS | NS | NS | NS | NS | NS | * | * | * | * | * | * | * | * | * | * |
| 60 | NS | NS | NS | NS | NS | NS | NS | NS | NS | NS | * | * | * | * | * | NS | NS | NS | NS | NS |
| 70 | NS | NS | NS | NS | NS | NS | NS | NS | NS | NS | * | * | * | * | * | NS | NS | NS | NS | NS |
| 80 | NS | NS | NS | NS | NS | NS | NS | NS | NS | NS | * | * | * | * | * | NS | NS | NS | NS | NS |
| 90 | NS | NS | NS | NS | NS | NS | NS | NS | NS | NS | * | * | * | * | * | NS | NS | NS | NS | NS |
| 100 | NS | NS | NS | NS | NS | NS | NS | NS | NS | NS | * | * | * | * | * | NS | NS | NS | NS | NS |

**Table S5:**
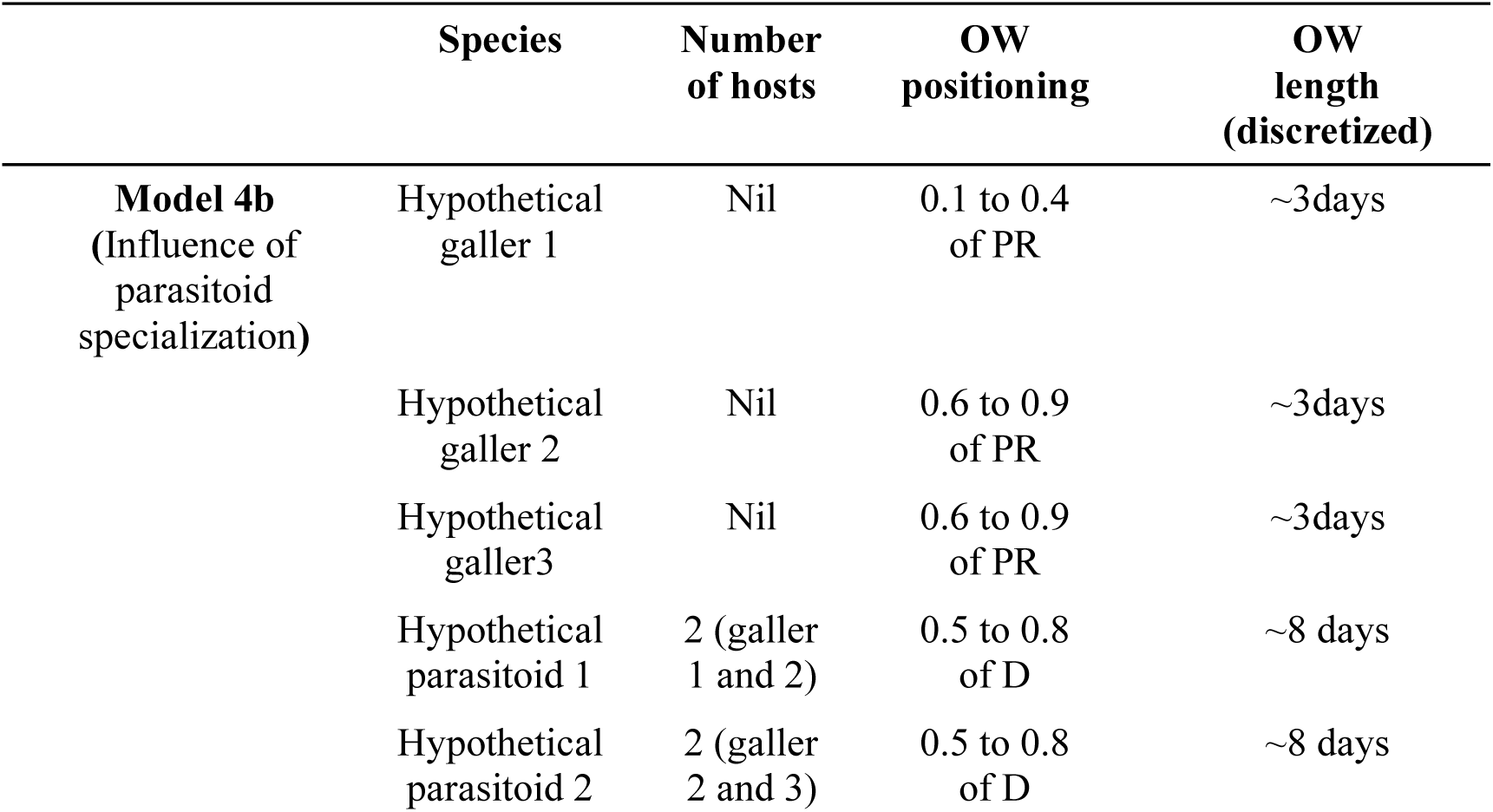
Details for variant of Model 4 (to test if the additive length of the oviposition windows of prey influence parasitoid colonization success).

